# SMAD4 governs a feedforward regulation of the TGF-β-effects in CD8 T cells that contributes to preventing chronic intestinal inflammation

**DOI:** 10.1101/2021.02.19.431987

**Authors:** Ramdane Igalouzene, Hector Hernandez-Vargas, Nicolas Benech, David Bauché, Célia Barrachina, Emeric Dubois, Julien C. Marie, Saïdi M’Homa Soudja

## Abstract

SMAD4, a key mediator of TGF-β signaling, plays a crucial role in T cells to prevent chronic gut inflammation. However, the molecular mechanisms underlying this control remain elusive. Using different genetic and epigenetic approaches, we unexpectedly reveal that SMAD4 in CD8 T cells prevents chronic intestinal inflammation by a feedforward mechanism that is TGF-β-independent. Prior to any TGF-β-receptor engagement, SMAD4 acts as an active and basal repressor of epigenetic, transcriptional and functional TGF-β imprinting in CD8 T cells. Thus, in sharp opposition to total TGF-β signaling deletion, SMAD4 deletion impairs naïve CD8 T cell effector predisposition but promotes CD8 T cell accumulation and epithelial retention by promoting their response to IL-7 and their expression of integrins such as *Itgae*. Besides, SMAD4 deletion unleashes the induction of a wide range of TGF-β-signaling-repressors such as *Smad7, Ski, Skil*, and *Smurf2* and hampers TGF-β-mediated CD8 T cell immunosuppression. Mechanistically, prior to any TGF-β signal, SMAD4 binds to the loci of several TGF-β-target genes, and by regulating histone acetylation, represses their expression. The massive gut epithelial colonization, associated with their escape from the immunoregulatory TGF-β effects overtakes their poor effector preconditioning and elicits microbiota-driven chronic epithelial CD8 T cell activation. Hence, in an anticipatory manner, independently of TGF-β, SMAD4 governs a feedforward regulation of TGF-β effects in CD8 T cells, preventing chronic intestinal inflammation.

## Introduction

An excessive immune reaction against microbiota is widely regarded as a common feature of inflammatory bowel diseases (IBDs) ^1,2^. Several immune-regulatory mechanisms prevent this reaction in the gastrointestinal tract, including the presence of the transforming growth factor beta (TGF-β) cytokine. This highly conserved cytokine is abundantly produced in the mammalian gut, ^3^ and is strongly implicated in immune cell regulation, and particularly T lymphocyte regulation by repressing numerous effector T cell functions ^4–6^ and promoting regulatory T cell-development, stability and function ^7^.

The active form of TGF-β binds to TGF-βRII leading to its auto-phosphorylation, which in turn phosphorylates TGF-βRI through its kinase domain. TGF-βRI then induces the phosphorylation of SMAD2 and SMAD3, which subsequently interact with either SMAD4 or tripartite motif-containing 33 (TRIM33). These complexes translocate to the nucleus and regulate the expression of several gene sets depending on the cellular and molecular context, including a wide range of TGF-β-repressor genes such as *Smad7* and *Ski*, thus generating a negative feedback loop to finely control TGF-β signaling ^8–10^. In addition, other non-canonical signaling pathways have been described downstream of TGF-βR engagement, namely MAPK/MEK, JNK/p38 and AKT/PI3K ^11^.

The signaling pathways activated by TGF-β can work either in concert or in opposition depending on the context, hampering their deciphering. For instance, during hematopoiesis each pathway regulates distinct and selective sets of genes in response to TGF-β, thus regulating different steps of hematopoiesis ^12^. Aside from this complementary interplay, TGF-β-activated pathways can also compete and inhibit each other. For instance, TRIM33 can compete with SMAD4 to interact with SMAD2 and SMAD3, and in addition, can induce SMAD4 degradation through its E3 ubiquitin ligase function ^13–15^. Given this intricate interplay, ablating one branch of TGF-β signaling may functionally impact the others, depending on the context.

TGF-β signaling is altered in IBD and CRC patients ^16^. Indeed, even though the gut mucosa of those patients displays a high level of TGF-β ^17^, their intestinal T cells are poorly responsive to this anti-inflammatory cytokine owing to the overexpression of the TGF-β repressor, SMAD7. In addition, patients harboring SMAD4 germline mutations develop intestinal polyps and are more predisposed to develop CRCs ^18^. This has further been demonstrated using genetic mouse models, in which SMAD4 deficiency in T cells drove chronic inflammation and cancer, highlighting a crucial role for SMAD4 in T cells in preventing IBDs ^19–21^. However, the precise cellular and molecular mechanisms governed by SMAD4 in directing this protective role remain undetermined.

Here, we reveal that SMAD4, in an anticipatory manner, independently of TGF-β, orchestrates a feedforward control of TGF-β-effects in CD8 T cells that is crucial in preventing IBDs. Consequently, SMAD4 ablation endows CD8 T cells with a strong epigenetic, -transcriptional and -functional TGF-β signature. This TGF-β-independent SMAD4 function restricts the accumulation and the intestinal epithelial retention of CD8 T cells. Besides, we uncover that SMAD4, before any TGF-βR-engagement, directly inhibits TGF-β negative feedback loop mediator expression. Thus, by reducing the basal TGF-β target gene and TGF-β repressor expression, SMAD4 potentiates to TGF-β-effect. Mechanistically, prior to any TGF-β signaling, SMAD4 acts at the chromatin level and regulates numerous TGF-β target genes by epigenetic modifications inversely to TGF-β signaling. Altogether, our findings unveil an original SMAD4 feedforward regulation in CD8 T cells that predisposes CD8 T cell to TGF-β-effects and contributes grandly to maintain intestinal homeostasis.

## Results

### SMAD4 in T cells protects mice from IBDs in a TGF-β-independent manner

Given the intricate interplay between TGF-β pathways, we first investigated the impact of the other TGF-β branches in the gut inflammation described in mice lacking SMAD4 in T lymphocytes. To this end, we used the CD4-CRE conditional deletion system to establish several mouse strains lacking one or several TGF-β signaling branches. We established mice with a deletion of SMAD4 (SKO), TRIM33 (TKO), double deletion of TRIM33 and SMAD4 (STKO) or double deletion of TGF-βRII and SMAD4 (R2SKO) in T cells **(Fig. 1a, Supplementary information, Fig. 1a)**. Consistent with previous studies, TKO, SKO, STKO and R2SKO mice did not display any signs of autoimmunity even at a more advanced age ^22–24^. However, strikingly, the weight of all mice lacking SMAD4 in T cells (SKO, STKO and R2SKO) stopped increasing from 4 months of age onwards **(Fig. 1b).** Postmortem analysis revealed an important intestinal inflammation in these animals, illustrated by an enlargement of the duodenum and a shortening of the colon **(Fig. 1c-d)**. Histological analysis revealed massive immune cell infiltrations in the mucosa and submucosa in both the small intestine and the colon of these mice compared to WT and TKO mice. Additionally, evident hyperplasia, crypt abscesses and strong intestinal crypt inflammation, likely cryptitis, were detected in all mice lacking SMAD4 in T cells, indicative of a strong chronic inflammation in these animals **(Fig. 1e, Supplementary information, Fig.2b)**. Collectively, our data genetically demonstrate that the ablation of the remaining TGF-β pathways in SKO mice does not prevent mice from developing chronic intestinal inflammation.

**Figure 1:**
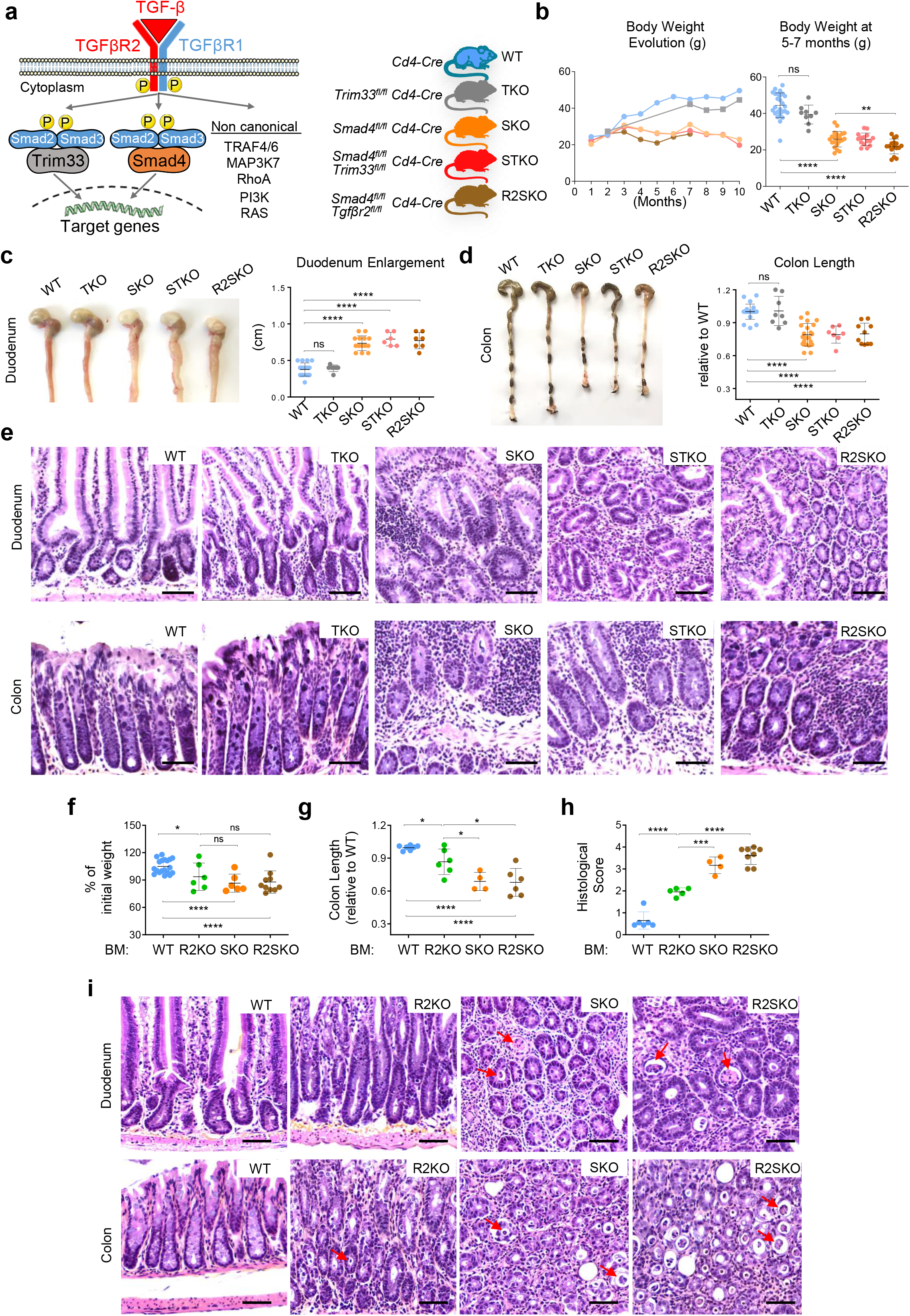
SMAD4 in T cells prevents chronic intestinal inflammation largely in a TGF-β - independent manner. **(a):** Scheme representing pathways of TGF-β signaling and mice models. **(b):** On the left panel, body weight of mice from 1 to 10 months old (n= 2 to 10 mice per group for each time point) and on the right panel, weight of the mice at 5-7 months old (n=6 to 10 mice per group). All mice are male. **(c-d):** Representative pictures of colon and duodenum, colon length and duodenum enlargement of the different strains of mice at 7 months of age. **(e):** Representative Hematoxylin & Eosin (H&E) staining of duodenum and colon sections of different mouse strains at 7 months old. Scale bar represents 50μm. **(f-i):** Irradiated RAG2KO mice were reconstituted with WT, R2KO, SKO, or R2SKO bone marrow (BM) cells; Percentage change in body weight between the beginning and the end of experiment **(f)**; colon length **(g)**; histological intestinal damage score **(h)**; and representative Hematoxylin & Eosin (H&E) staining of duodenum and colon sections **(i).** Scale bar represents 50μm.Red arrows highlight crypt abscesses. All Data represent at least 3 independent experiments (C, D, E, F, G, H, I) and presented as mean ± SD. Each symbol represents an individual mouse. Data were analyzed by unpaired Student t Test. ns: nonsignificant; * p<0,05; **p < 0.01; ***p < 0.001; ****p < 0.0001.

Then, given that SMAD4 is known to mediate key biological functions independently of TGF-β signaling in T cells ^22,25^, we assessed whether a TGF-β-independent SMAD4 function in T cells could contribute to maintain intestinal homeostasis. Given that TGF-βRII-deficient (R2KO) mice die within 3-4 weeks ^5,6^ and intestinal inflammation only develops at 5 months in SMAD4-deficient mice, we employed a bone marrow (BM)-engrafted mouse model to compare age-matched adult mice. We engrafted irradiated adult mice with BM from WT, R2KO, SKO, and R2SKO mice. Mice engrafted with R2KO, SKO, and R2SKO BM cells lose weight compared to WT engrafted ones **(Fig. 1f)**. However, mice engrafted with BM cells from SKO and R2SKO mice developed more severe gut inflammation compared to those engrafted with R2KO BM cells, as evidenced by shorter colons, massive immune cell infiltrations, hyperplasia and important mucosal damage **(Fig. 1g-i, Supplementary information, Fig.1c)**. Hence, these observations demonstrate that SMAD4 in T cells, in a TGF-β independent manner, contributes to maintain intestinal homeostasis.

### CD8αβ T cells are key effector cells that contribute to the intestinal immunopathology observed in SMAD4 deficient mice

Next, we examined which effector T cell population mediates this intestinal immunopathology by using specific anti-CD4 and anti-CD8β depleting antibodies. To avoid undesirable long-term side effects of depleting antibody treatment, we used the BM-engrafted mouse models **(Fig. 2a)**. Flow cytometry analysis confirmed the effective ablation of conventional CD8αβ and CD4 T cells in secondary lymphoid organs (SLOs) and in the gut without depleting the other populations, such as CD8αα TCRαβ and TCRγδ populations **(Supplementary information, Fig. 2a-b)**. Remarkably, BM SKO-engrafted mice treated with anti-CD8β did not exhibit weight loss and colon length reduction, in sharp contrast to anti-CD4 treated mice **(Fig. 2b-c)**. Furthermore, histological examination showed a substantial decrease in immune cell infiltration and absence of hyperplasia and crypt abscesses in BM SKO-engrafted mice treated with anti-CD8β **(Fig. 2d)**. Similarly, anti-CD8β treatment rescued BM R2SKO-engrafted mice from developing intestinal inflammation **(Supplementary information, Fig. 2c-d)**. Collectively, these results suggest a key effector role of CD8αβ T cell in contributing to the intestinal damage in SMAD4 deficient mice.

**Figure 2:**
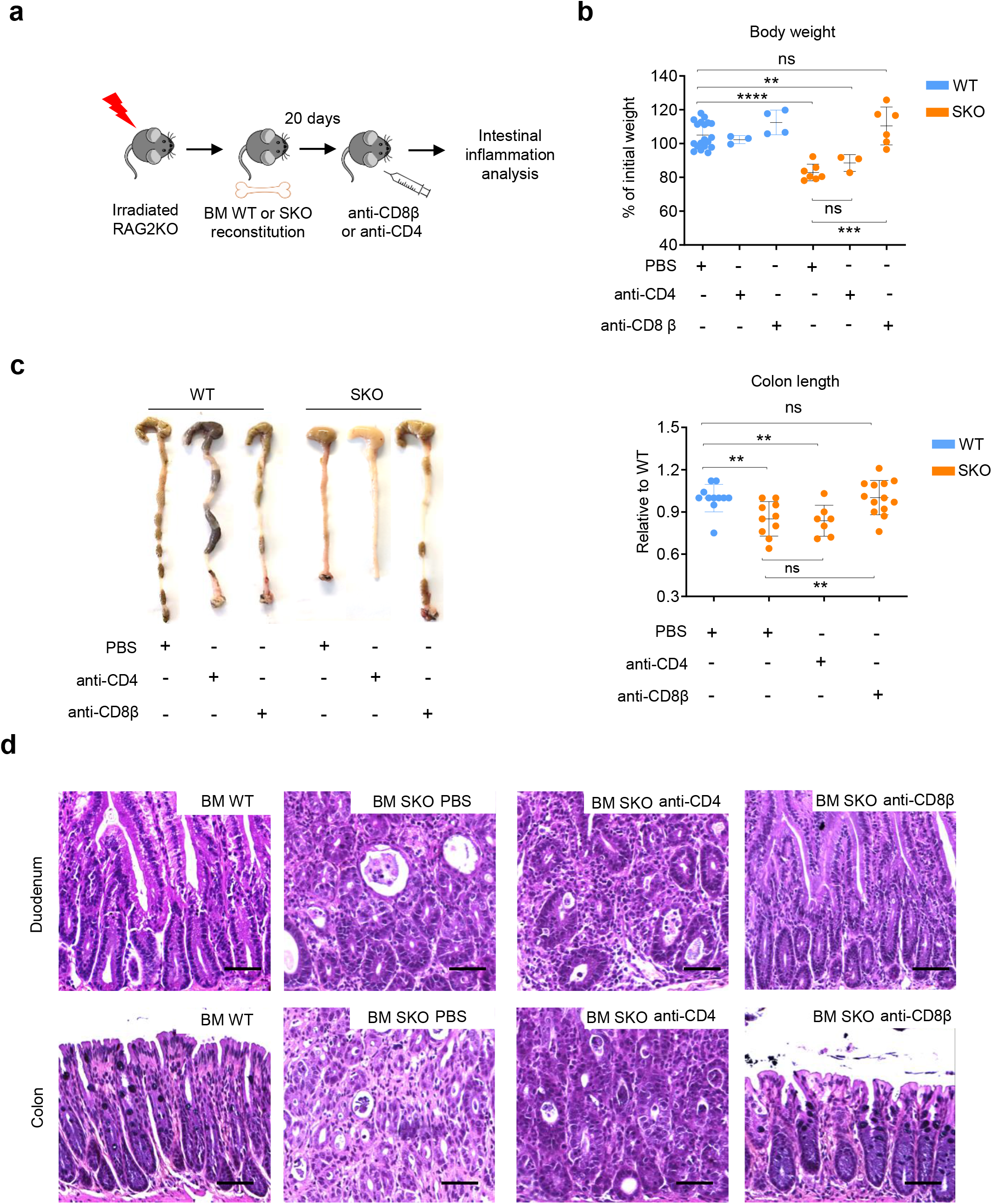
CD8αβ T cell-depletion prevents intestinal inflammation upon SMAD4 deletion in T cells. **(a):** Scheme of the *in vivo* CD8β and CD4 depletion experiment. RAG2KO mice were sub-lethally irradiated and reconstituted with WT or SKO BM cells. 20 days after reconstitution, mice were injected or not intraperitoneally with an anti-CD8β or anti-CD4 depleting antibody. **(b):** Body weight at day 40 after WT or SKO BM reconstitution and treatment with anti-CD8β or anti-CD4 depleting antibody. **(c):** Representative pictures of colons and colon length measurement of BM reconstituted mice, treated with anti CD8β or anti-CD4 depleting antibody. **(d):** Representative Hematoxylin & Eosin (H&E) staining of duodenum and colon sections of irradiated mice reconstituted with WT or SKO BM cells and treated with anti-CD8β or anti-CD4 depleting antibody. Scale bar represents 50μm. All data represent at least 3 independent experiments and presented as mean ± SD. Each symbol represents an individual mouse. Data were analyzed by unpaired Student t test. ns: nonsignificant; * p<0,05; **p < 0.01; ***p < 0.001; ****p < 0.0001.

### SMAD4 prevents microbiota-driven accumulation and activation of CD8αβ T cells within the gut-epithelium

We then assessed the mechanism by which SMAD4 in CD8αβ T cells prevents intestinal immunopathology. Strikingly, we observed in all SMAD4-deficient mice (SKO, STKO and R2SKO) a substantial increase in the frequency and numbers of CD8αβ T cells in secondary lymphoid organs, as well as in the lungs, skin, colon, and small intestine, compared to WT or TKO mice **(Fig. 3a-b, Supplementary information, Fig. 3a-b).** This data revealed a systemic accumulation of CD8αβ T cells in the absence of SMAD4. Besides this important accumulation, CD8αβ T cells from SKO, STKO and R2SKO mice, expressed large amounts of cytotoxic molecules including granzymes A and B (GZMA and GZMB) or pro-inflammatory cytokines and chemokines such as IFN-γ, TNFα and CCL3 in the intestinal epithelium compared to WT and TKO mice **(Fig. 3c, Supplementary information, Fig. 3c-d)**. Importantly, the strong co-expression of the epithelial retention marker CD103 and GZMB, suggests that the activated CD8αβ T cells were likely *bona fide* intra-epithelial lymphocytes (IELs) **(Supplementary information, Fig. 3e)**. Remarkably, CD8αβ T cells from SMAD4-deficient mice were barely or not activated in the spleen, the lung, skin, lymph nodes, and *lamina propria* of the intestine **(Fig. 3c, Supplementary information, Fig. 3f-g)**. This indicated a spatial-restricted-activation of SMAD4-deficient CD8 T cells within the intestine epithelium.

**Figure 3:**
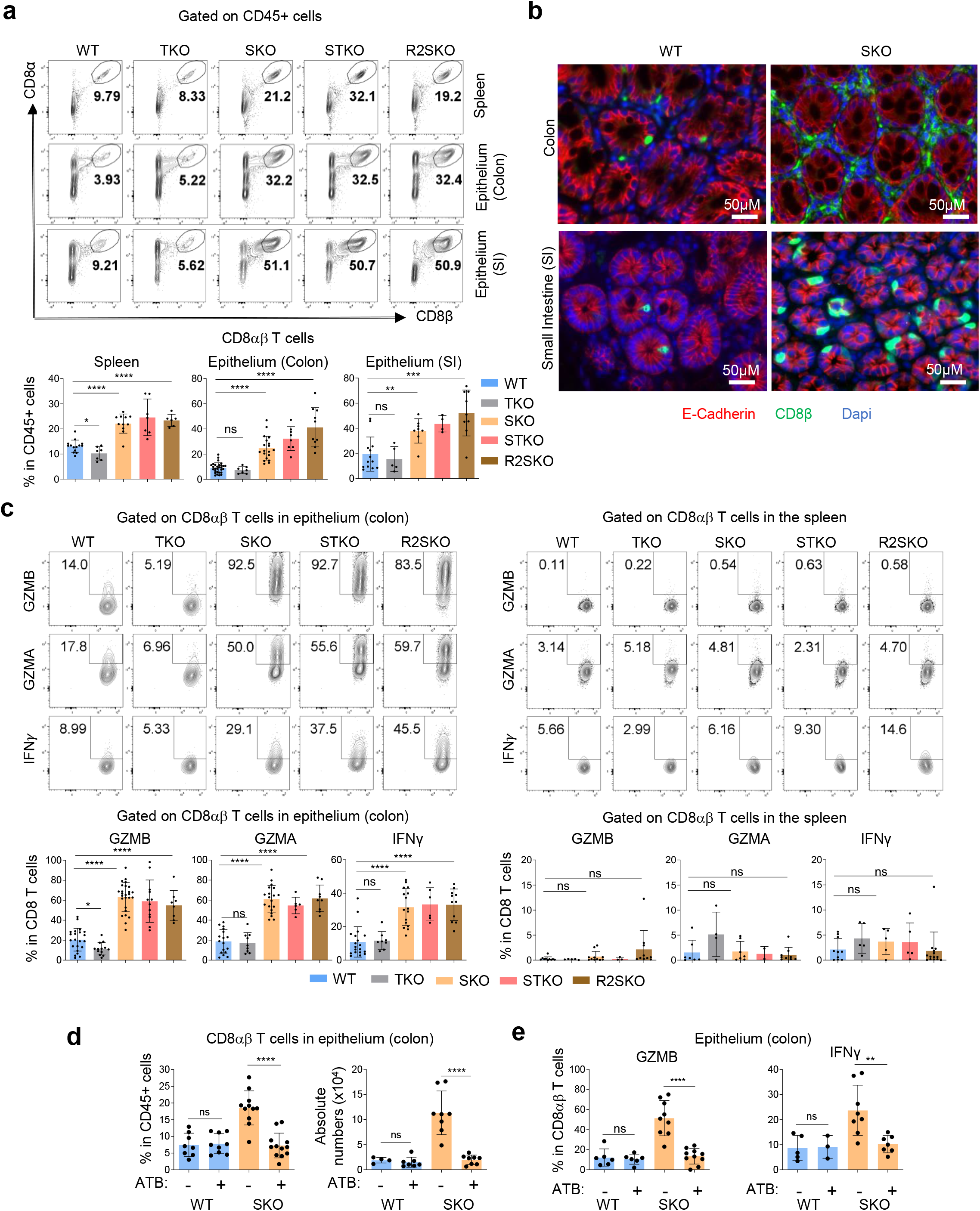
SMAD4 in T cells prevents microbiota-mediated accumulation and epithelial activation of CD8αβ T cells. **(a):** Representative flow cytometry data showing the frequency of CD8αβ T cells among CD45+ cells in the spleen and epithelium from the colon and small intestine of 7 months aged mice, (n= minimum 4 mice per group). **(b):** Representative pictures showing immune-fluorescence staining of CD8β (green), E-cadherin (red), DAPI (blue) in the small intestine and colon sections of 7 months-aged WT and SKO mice. **(c):** Flow cytometry staining of GZMA, GZMB and IFN-γ among splenic and colonic intra-epithelial lymphocytes CD8αβ T cells. **(d-e):** Effect of antibiotic (ATB) treatment on the frequency, numbers, and activation of colonic intraepithelial CD8αβ T cells from WT and SKO mice. All data represent at least 3 independent experiments and presented as mean ± SD. Each symbol represents an individual mouse. Data were analyzed by unpaired Student t test. ns: nonsignificant; * p<0,05; **p < 0.01; ***p < 0.001; ****p < 0.0001.

Next, we investigated the mechanisms triggering intestinal epithelial activation of CD8αβ T cells in SMAD4-deficient mice. Given the importance of the microbiota in shaping intestinal immunity and promoting IBDs ^1,2^, we hypothesized that commensal bacteria could be responsible for CD8αβ T cell intestinal epithelial accumulation and exacerbated epithelial activation. In order to confirm this scenario, SKO mice were treated with antibiotics (ATB). Strikingly, ATB treatment of SKO mice completely abrogated CD8αβ T cell accumulation in the gut epithelium **(Fig. 3d)**. In addition, the enhanced production of IFN-γ and granzymes in CD8αβ IELs was also abolished in ATB-treated SKO mice **(Fig. 3e)**. Hence, these data reveal that the TGF-β-independent SMAD4 function prevents the spontaneous microbiota-driven activation of CD8αβ T cells within the epithelial layer of the intestine.

### In the absence of TGF-β, SMAD4 restrains the TGF-β-transcriptional signature in CD8 T cells and preconditions effector differentiation of naïve CD8 T cells

To go deeper in the molecular processes governing SMAD4-mediated control of intestinal homeostasis in CD8 T cells, we next performed a global gene expression profile of CD8 T cells from WT, SKO and R2KO mice. In order to rule out any potential side effects of the inflammatory environment, we used F5 TCR transgenic CD8 T cells in a RAG2KO background, since these mice did not develop any inflammation **(Supplementary information, Fig. 4a)**. Unexpectedly, the comparison between SKO CD8 T cells and R2KO CD8 T cells resulted in a larger set of significantly differentially expressed genes (DEGs) (1573 genes) (FDR <0.05) than the comparison between SKO and WT mice (408 DEGs) **(Fig. 4a)**, highlighting a wider molecular gap between SKO and R2KO naïve CD8 T cells. An unsupervised hierarchical clustering of all DEGs revealed five distinct clusters. Strikingly, DEGs in which the SMAD4 deletion and the TGF-βRII deletion show a distinct expression pattern (clusters II, III and V) represent more than 92 % of all DEGs, definitively, indicating a wide transcriptional disparity between SKO and R2KO CD8 T cells. **(Fig. 4b, Supplementary information, Fig. 4b)**. More importantly, the large majority of the divergent DEGs are genes where SMAD4 deletion affects negatively (cluster II) or positively (cluster III) their expression compared to WT and oppositely to TGF-β signaling depletion (**Fig. 4b-c**). Thus, this wide transcriptional disparity is largely attributed to an evident opposition between SMAD4 and TGF-β signaling and reveals that SMAD4 acts as a repressor of TGF-β transcriptional outcome in CD8 T cells. To assess whether this marked transcriptional opposition orchestrated by SMAD4 is not mediated by a TGF-β signal, we then conducted a genome-wide RNA sequencing in R2SKO CD8 T cells and compared the gene expression profiles of CD8 T cells from F5 R2SKO with F5 SKO or F5 R2KO mice. This analysis unveiled a larger set of DEGs (740 genes) in the comparison between R2SKO and R2KO CD8 T cells by contrast to the comparison between SKO and R2SKO CD8 T cells (106 DEGs) **(Fig. 4d)**. Furthermore, the absence of SMAD4 largely reverts the gene overexpression (and gene downregulation) observed after total TGF-β signaling deletion in R2SKO CD8 T cells **(Fig. 4e).** Thus, SMAD4, in the absence of TGF-β, in an anticipatory manner, acts as a basal and active repressor of TGF-β-transcriptional landscape in CD8 T cells.

**Figure 4:**
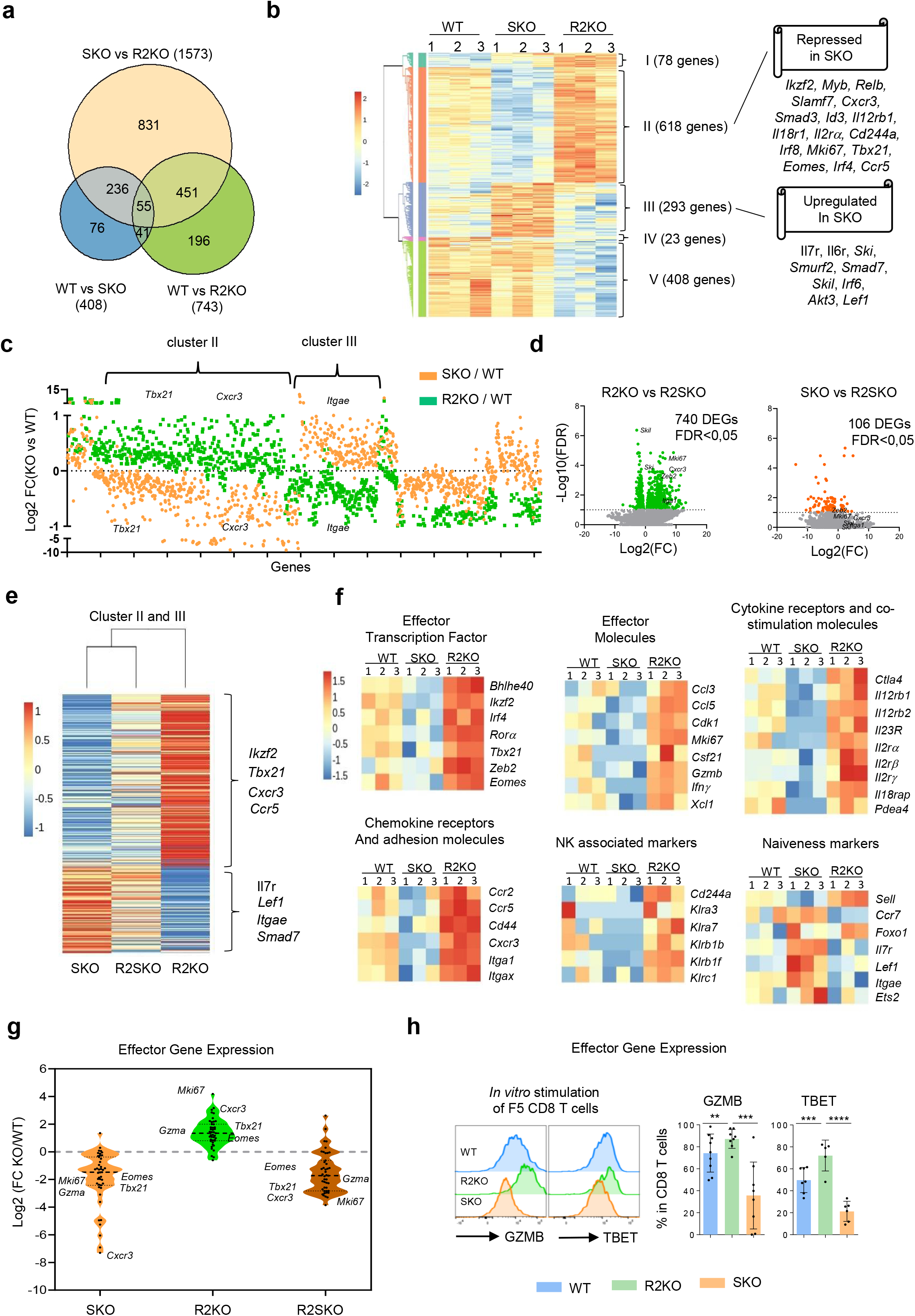
In absence of TGF-β signal, SMAD4 represses TGF-β signature in naïve CD8αβ T cells inversely of TGFβRII signaling. **(a):** Venn diagram showing the numbers of differentially expressed genes between WT, R2KO, and SKO naïve F5 CD8αβ T cells. **(b):** Heatmap showing the hierarchical clustering of differentially expressed genes between WT, SKO, and R2KO F5 naïve CD8αβ T cells. **(c):** Fold change (logarithmic scale) of gene expression of SKO vs WT (in orange) and R2KO vs WT (in green). DEGs correspond to those shown in heatmap 4B **(d):** Volcano plot of RNA-seq data from R2KO, SKO, and R2SKO naïve F5 CD8αβ T cells. The data for all genes is plotted as log2 fold change versus the -log10 of the adjusted p-value. Genes selected as significantly different are highlighted as green and red dots. Some example genes are labelled with gene symbols**. (e):** Heatmap showing the Log2 fold change expression of genes of cluster II and III highlighted in Fig.2b, and for each condition, the heatmap value corresponds to the KO relative to WT (average of 3 biological replicates). **(f):** Heatmaps showing the expression of genes linked to CD8 T cell effector functions and genes linked to naïve and quiescence stage on WT, SKO or R2KO F5 naïve CD8αβ T cells. **(g):** Violin plot showing the relative expression of effector genes from R2KO, SKO, and R2SKO CD8 T cells compared to WT**. (h):** Flow cytometry staining for GZMB and TBET in F5 CD8αβ T cells after anti CD3/CD28 stimulation for 4 days *in vitro*. All data represent at least 3 independent experiments and presented as mean ± SD. Each symbol represents an individual mouse. Data were analyzed by unpaired Student t test. ns: nonsignificant; * p<0,05; **p < 0.01; ***p < 0.001; ****p < 0.0001.

Then we determined functional outcomes of this impressive transcriptional divergence. A deeper examination of the divergent DEGs highlights many genes belonging to the T cell effector program. In SMAD4-deficient naïve CD8 T cells, genes encoding effector molecules such as *Ifnγ IL12rβ2, Gzmb, Gzma, Gzmk*, and *Cd244* were repressed. Accordingly, the expression of transcription factors known to direct CD8 T cell effector differentiation (*Tbx21, Irf4, Zeb2*, and *Eomes*) was also attenuated **(Fig. 4f)**. Conversely, genes associated with quiescence/naiveness of CD8 T cells (eg. *Lef1, itgae, Il7r, Ets2*) were slightly enhanced or not significantly affected in SMAD4-deficient CD8αβ T cells. Single TGF-βRII deletion, in contrast, drastically promoted the expression of effector genes **(Fig. 4f)**. A gene set enrichment analysis (GSEA) of all DEGs and the expression of 43 selected genes associated with T cell activation indicated that similarly to SKO CD8 T cells, the effector gene predisposition was also repressed in naïve R2SKO CD8 T cells **(Fig. 4g, Supplementary information, Fig. 4c-d)**. Functionally, CD8 T cells lacking SMAD4 displayed less activation judged by the weaker GZMB and TBET expression compared to WT and R2KO cells after *in vitro* activation **(Fig. 4h)**. Overall, our data reveal that in the absence of TGF-β, SMAD4 restricts transcriptional and functional TGF-β signature in CD8 T cells and endows naïve CD8 T cells with an effector program.

### SMAD4 facilitates CD8 T cell response to TGF-β, in a TGF-β-independent manner

Since SMAD4 deletion limits T cell activation, a compensatory mechanism must allow microbiota-driven activation of CD8 T cells in the gut. Intriguingly, genes encoding potent TGF-β signaling repressors (e.g. *Smad7, Ski, Skil* and *Smurf2*) were enhanced in SKO and R2SKO compared to R2KO CD8 T cells **(Fig. 5a)**. We validated the overexpression of those genes by real-time quantitative RT-PCR on naïve F5 CD8 T cells, as well as on polyclonal CD8 T cells **(Fig. 5b, Supplementary information, Fig.5a)**, attesting that this overexpression was not restricted to a specific T cell receptor (TCR) repertoire. The expression defect of TGF-β repressors in R2KO CD8 T cells confirmed that they are TGF-β target genes ^8,9,26^. Since the double deletion of TGF-βRII and SMAD4 (R2SKO) restored the gene expression of TGF-β repressors **(Fig. 5b, Supplementary information, Fig.5a)**, this demonstrated definitively that SMAD4 inhibits the expression of TGF-β repressors in a TGF-β-independent manner.

**Figure 5:**
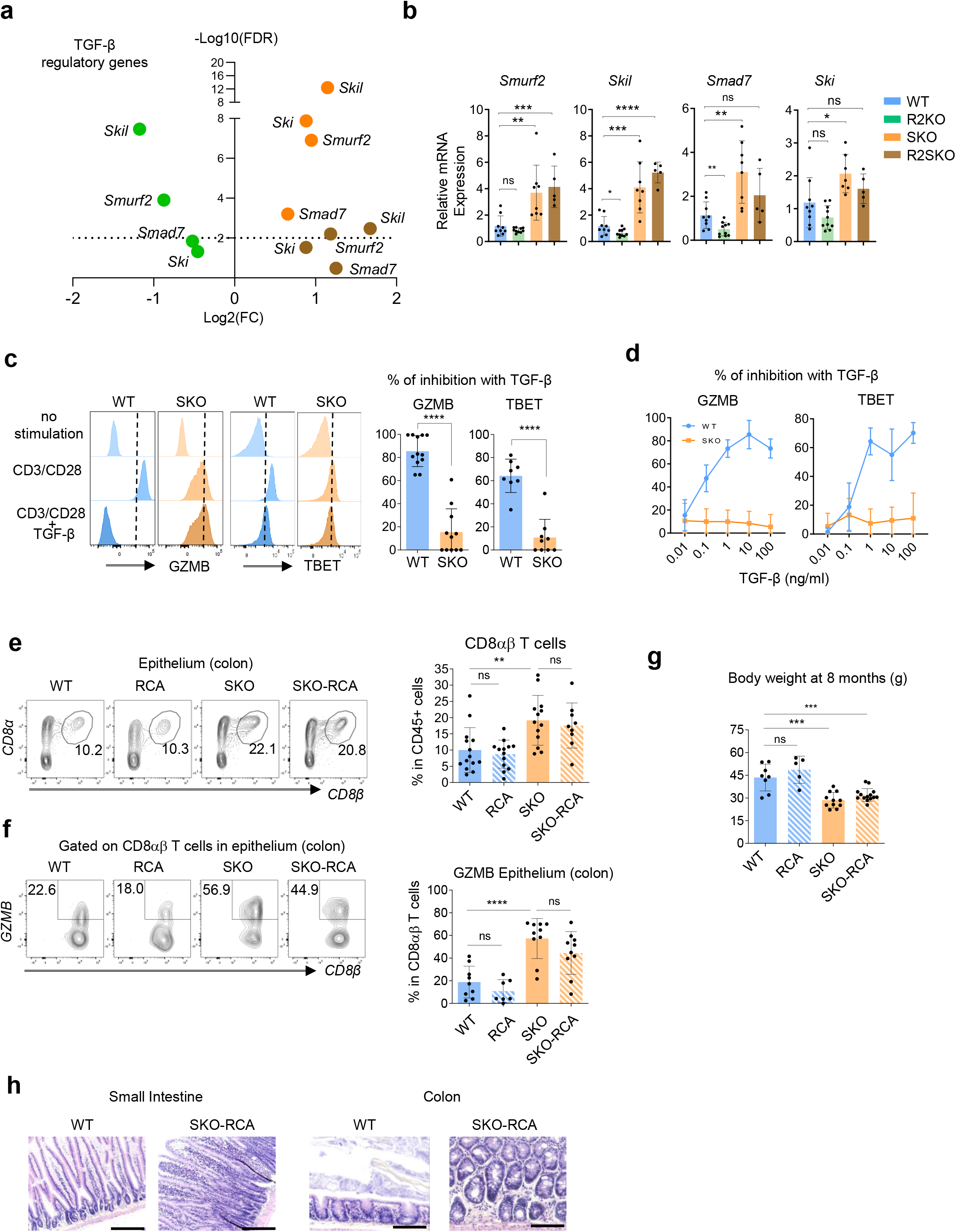
SMAD4 depletion promotes expression of TGF-β repressors and impedes TGF-β response in CD8αβ T cells *in vitro* and *in vivo*. **(a):** Volcano plot showing TGF-β inhibitory genes in SKO (orange), R2KO (green), and R2SKO (brown) F5 naïve CD8αβ T cells, all relative to WT. **(b):** Quantitative RT-PCR analysis of the expression of indicated TGF-β regulatory genes in F5 naïve CD8αβ T cells from spleen of WT, R2KO, SKO, and R2SKO mice (n=5-6). These mice are different from those used for RNA-seq data. **(c-d):** Flow cytometry data showing WT or SKO CD8αβ T cell-inhibition of GZMB and TBET after anti CD3/CD28 stimulation with or without recombinant TGF-β at 10ng/ml **(c)** or different concentrations **(d)**. The percentage of inhibition of CD8αβ T cells was appreciated by calculating the ratio between anti CD3/CD28 + TGF-β condition and anti CD3/CD28 alone. **(e-f):** representative flow cytometry plots showing the frequency of CD8αβ T cells among CD45+ cells present within the colonic epithelium **(e),** and intra cellular staining for GZMB among colonic epithelial CD8αβ T cells **(f);** Body weight **(g);** and Hematoxylin & Eosin (H&E) staining of duodenum and colon sections **(h)**; from 8 months aged WT, RCA, SKO and SKO-RCA mice, (n= minimum 6 mice per group). Scale bar represents 200μm. All data represent at least 3 independent experiments and presented as mean ± SD. Each symbol represents an individual mouse. Data were analyzed by unpaired Student t test. ns: nonsignificant; * p<0,05; **p < 0.01; ***p < 0.001; ****p < 0.0001.

Since those genes are potent repressors of TGF-β signaling and have been associated with a defect of T cell response to TGF-β in IBDs ^27^, we examined SMAD4-deficient CD8 T cell TGF-β-response. While TGF-β inhibited impressively GZMB and TBET expression in activated WT CD8 T cells even at low doses, impressively, their expression was maintained even at high concentrations of TGF-β in SKO CD8 T cells **(Fig. 5c-d)**. Thus, these observations strongly reveal that SMAD4 ablation totally limits the immune-regulatory effects of TGF-β on CD8 T cells. Importantly, this demonstrates that SMAD4 is crucial for TGF-β-mediated immunosuppression and is not redundant. Because TGF-β is highly enriched in the gut ^3^ and represses T cell activation ^28^, this impaired response to TGF-β could contribute to the chronic microbiota-driven CD8 T cell activation. In order to confirm this assumption *in vivo*, we forced the activation of SMAD4 independent pathways of TGF-β signaling by crossing SKO mice with mice bearing a conditionally-expressed, constitutively-active form of the TGF-βR1 (LSL-TGFβRICA mouse strain) ^29^. In the resulting SKO-RCA mice animals, CD8αβ T cells were as abundant and activated in the gut epithelium as in SKO mice **(Fig. 5f-g)**, and more importantly, SKO-RCA mice developed IBDs **(Fig. 5h-i).** Hence, the remaining TGF-β signaling pathways are unable to compensate for SMAD4 loss. Collectively, these data suggest that the TGF-β-independent function of SMAD4 facilitates the response of CD8 T cells to TGF-β, by restraining the expression of a wide range of TGF-β repressors in a feedforward mechanism (prior to any TGF-β signal) and this is crucial and non-redundant to mediate immune-regulatory-effect of TGF-β.

### SMAD4 restrains homeostatic survival and epithelial retention of CD8 T cells in a TGF-β-independent manner

Given that R2KO mice in which T cells do not respond to TGF-β signal, do not exhibit gut inflammation as severe as in SKO and R2SKO mice **(Fig. 1f-i)**, additional factors might enhance the intestinal inflammation in SKO and R2SKO mice. Strategically, we focused on genes crucial for CD8 T cell homeostasis and epithelial layer retention that are similarly affected in SKO and R2SKO and inversely in R2KO CD8 T cells. Our first target was IL-7R, also termed CD127, since it plays a crucial and non-redundant role in homeostatic survival of CD8 T cells and recent studies associated IL-7 signaling over-activation and IBDs ^30–33^ In line with the RNA-seq data, flow cytometry analysis validated that naïve F5 SMAD4-deficient (SKO and R2SKO) CD8 T cells overexpressed IL-7R compared to WT CD8 T cells, in sharp contrast to R2KO CD8 T cells **(Fig. 6a)**. Similarly, we observed this upregulation in CD8 T cells with a polyclonal TCR repertoire **(Supplementary information, Fig. 6a)**. Consistent with the level of IL-7R expression, STAT5 phosphorylation, which is induced upon IL-7 stimulation, was slightly enhanced in SKO and R2SKO CD8 T cells, and impaired in R2KO CD8 T cells **(Fig. 6b)**. A time-course analysis of survival demonstrated that IL-7 did not prevent R2KO CD8 T cells from dying *in vitro* compared to SKO and R2SKO CD8 T cells that survived largely better **(Fig. 6c)**. Accordingly, we observed a substantial increase in the absolute number and the proportion of CD8 T cells in secondary lymphoid organs from SKO and R2SKO F5 transgenic mice, unlike R2KO mice **(Fig. 6d and data not shown)**. These findings reveal a critical role for the TGF-β-independent SMAD4 function in restraining CD8 T cell accumulation by repressing the IL-7 response, in sharp contrast to TGF-β signaling.

**Figure 6:**
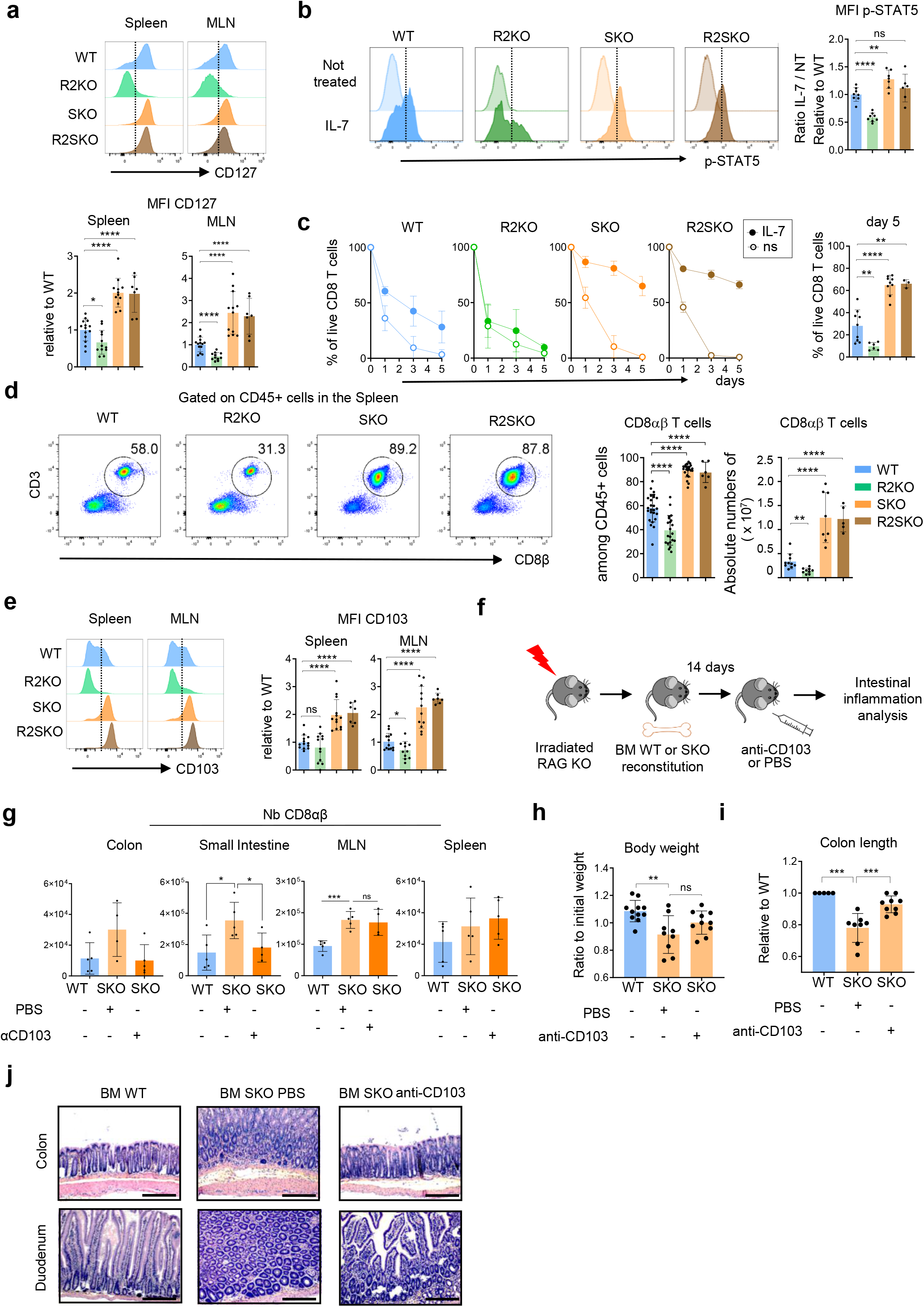
SMAD4 promotes homeostatic survival and epithelial retention of CD8αβ T cells in an opposite way to TGFβRII signaling. **(a):** Flow cytometry staining of CD127 on WT, R2KO, SKO and R2SKO F5 naïve CD8αβ T cells. **(b):** Flow cytometry staining of p-STAT5 after IL-7 *in vitro* treatment in WT, R2KO, SKO and R2SKO F5 naïve CD8αβ T cells. **(c):** Survival monitoring of WT, R2KO, SKO, or R2SKO naïve F5 CD8αβ T cells treated or not with IL-7. **(d):** Flow cytometry data showing the frequency with absolute numbers of F5 naïve CD8αβ T cells among CD45+ cells in the spleen of 3 months aged WT, R2KO, SKO and R2SKO F5 mice. **(e):** Flow cytometry staining of CD103 on WT, R2KO, SKO and R2SKO F5 naïve CD8αβ T cells. **(f-g):** Experimental procedure for anti-CD103 blocking treatment **(f)**; CD8 T cell numbers **(g),** Body weight **(h)**; colon length **(i)**; and Hematoxylin & Eosin (H&E) staining of duodenum and colon sections **(j)** of irradiated mice reconstituted with WT or SKO BM cells and treated or not with anti-CD103 blocking antibody. Scale bar represents 200μm. All data represent at least 3 independent experiments and presented as mean ± SD. Each symbol represents an individual mouse. Data were analyzed by unpaired Student t test. ns: nonsignificant; * p<0,05; **p < 0.01; ***p < 0.001; ****p < 0.0001.

Aside from *Il7r, Itgae* encoding for CD103 was also aberrantly upregulated in CD8 T cells from SKO mice. CD103 is of great interest as it elicits T cell retention within the intestinal epithelial layer ^34,35^. In agreement with the RNA-seq data, SKO and R2SKO naïve CD8 T cells exhibited an enhanced level of CD103 **(Fig. 6e)**. Similarly, we observed this upregulation in a polyclonal TCR repertoire **(Supplementary information, Fig. 6b)**. In correlation with the absence of CD103 expression, R2KO CD8 T cells are less enriched in the intestinal epithelium compared to R2SKO and SKO CD8 T cells **(Supplementary information, Fig. 6c, d and e)**. This impaired epithelial tropism of R2KO CD8 T cells may explain the milder intestinal inflammation observed on those mice compared to R2SKO mice. In line with this assumption, we next addressed whether the exacerbated expression of CD103 plays a role in the IBD observed in SKO mice. We treated BM-engrafted mice with a blocking antibody that specifically recognizes CD103 **(Fig. 6f)**. The CD103 blockade led to a decrease in CD8 T cell numbers within the intestinal epithelium of SKO mice, without altering their accumulation in secondary lymphoid organs such as the spleen and mesenteric lymph nodes **(Fig. 6g)**. Although this treatment did not fully restore body weight in SKO, the colon length and immune-histology analysis highlighted clear improvement **(Fig. 6h-j)**. The colon length reduction and the mucosal damage due to immune infiltration were alleviated, indicating a beneficial effect of CD103 blockade in SKO mice. Globally, in addition to the impaired response to TGF-β immune-regulatory functions, SMAD4 disruption promotes IL-7 responsiveness and epithelial retention of CD8 T cells in a TGF-β-independent manner. These combined alterations contribute to the outnumbering and the positioning of CD8 T cells in the gut epithelium of SKO mice leading to severe chronic intestinal inflammation compared to R2KO mice.

### SMAD4, in absence of TGF-β signal, binds to promoters and enhancers of a large set of TGF-β target genes to regulate their expression in a TGF-β-independent manner

To further decipher at the chromatin level the mechanisms by which SMAD4 regulates TGF-β signature imprinting in CD8 T cells, prior to any TGF-β signal, we conducted a chromatin immunoprecipitation sequencing (ChIP-seq) of SMAD4 on naïve CD8 T cells from WT, R2KO and SKO mice. 2982 peaks were identified in WT cells and 3432 peaks were identified in R2KO cells, demonstrating that SMAD4 binds to the genome even without TGF-β signal in CD8 T cells. Since most of the binding sites were localized in promoter regions (64% for the WT and 67% for the R2KO) or were closely located around the transcription start site (TSS) regions, this suggests grandly that SMAD4 directly regulates many variety of genes **(Fig. 7a-b, Supplementary information, Fig.7a)**. Accordingly SMAD4 binds irrespective or not to TGF-β signaling to genomic regions that regulate diverse biological pathways involved for instance in TCR signaling, RNA translation or TGF-β signaling regulation **(Supplementary information, Fig. 7b)**.This data highlights the broad potential impact of TGF-β-independent function of SMAD4 in diverse CD8 biological processes and emphases its interest. Then, we asked whether the genome-wide occupancy of SMAD4 is distinct with or without TGF-β signaling. Of the 2982 peaks in WT cells and 3432 peaks in R2KO cells, 1954 peaks were common, highlighting an important similarity in regional binding sites irrespective of the cellular response to TGF-β **(Fig. 7c).** Thus, this observation reveals that SMAD4, before any TGF-βR-engagement, occupies promoters and enhancers of different genes likely for regulating their expression.

**Figure 7:**
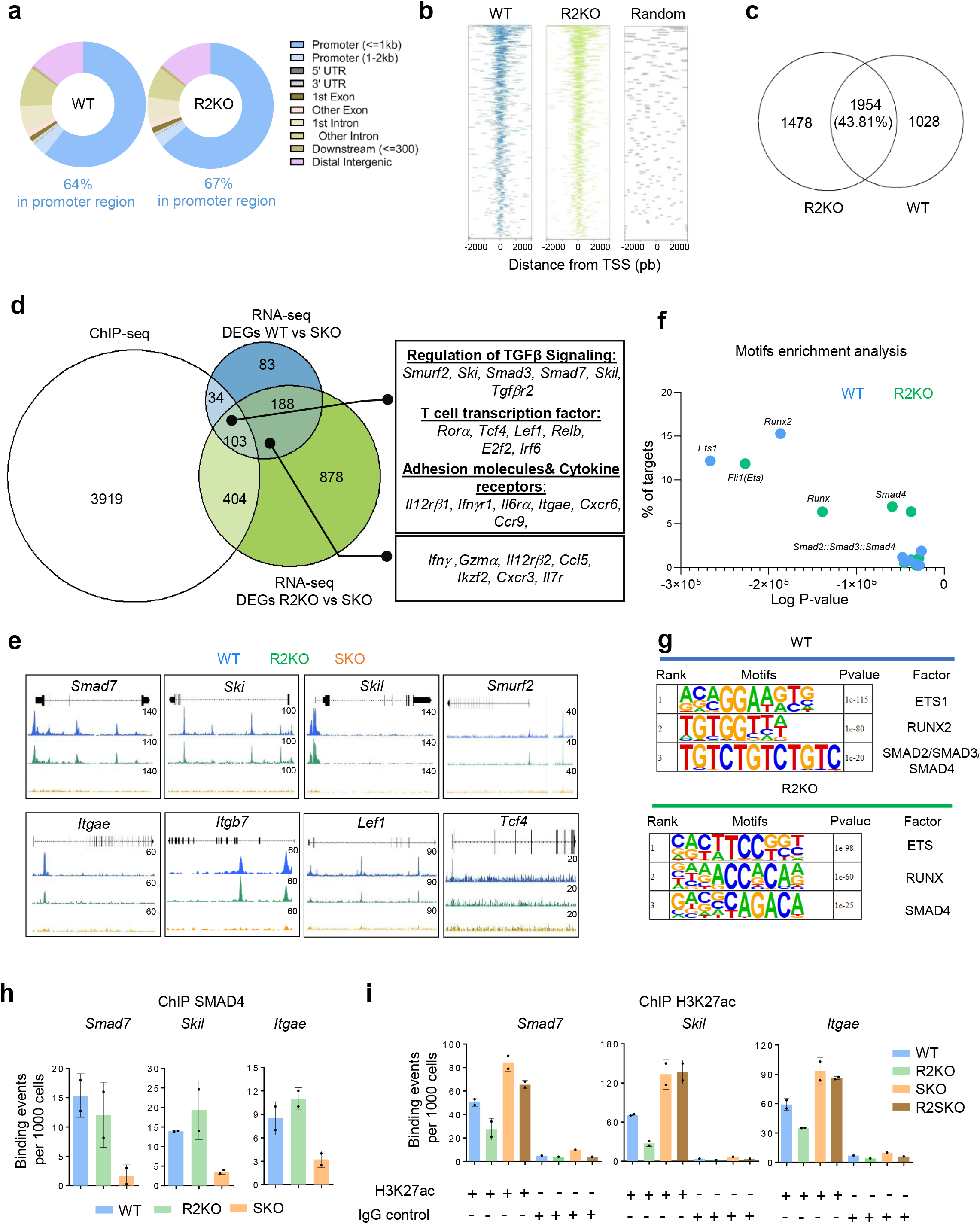
SMAD4 represses directly TGF-β target genes by histone deacetylation without requirement of TGF-β signaling. **(a):** The proportions of SMAD4 peaks associated with promoter, 5 UTR, 3 UTR, exon, intron, and intergenic regions in WT and R2KO naïve F5 CD8αβ T cells. **(b):** Enriched heatmaps showing the SMAD4-occupancy signals in genomically aggregated TSS regions in WT and R2KO CD8 T cells. Each panel represents 2 kb upstream and downstream of the TSSs **(c):** Venn diagram showing the number of SMAD4 common peaks between WT and R2KO naïve CD8αβ T cells. **(d):** Venn diagram showing the overlap between SMAD4 ChIP-seq peaks and RNA-seq DEGs. **(e):** SMAD4 binding ChIP-seq peaks in WT (blue), R2KO (green) or SKO control (orange), in corresponding genes. **(f):** Transcription factor (TF) top motifs in SMAD4 binding sites in WT and R2KO CD8 T cells. X-axis represents the logP-value of the motif enrichment. Y-axis represents the fold change of the motif enrichment**. (g):** The three top motifs found by Hypergeometric Optimization of Motif Enrichment (HOMER) analysis among SMAD4 binding peaks in WT and R2KO CD8 T cells. **(h):** q-PCR-based ChIP analysis of SMAD4 on the promoters/enhancers of *Smad7, Skil* and *Itgae* in WT, R2KO and SKO F5 naïve CD8αβ T cells. Each point represents a pool of minimum 3 mice. **(i):** qPCR-based ChIP analysis of H3K27ac on the promoters/enhancers of *Smad7, Skil* and *Itgae* in WT, R2KO, SKO, and R2SKO F5 naïve CD8αβ T cells. Each point represents a pool of minimum 3 mice.

By combining the ChIP-seq peaks of SMAD4 and the DEGs from RNA-seq data **(Figure 4)**, we found 541 genes that are potentially directly regulated by SMAD4 **(Fig. 7d)**. Focusing on genes that were differentially expressed between WT and SKO cells and between R2KO and SKO cells, we found 103 genes. Among those 103 genes, we found genes implicated in CD8 T cell differentiation such as *Tcf4* and *Lef1* but also many well-characterized TGF-β-target genes. Importantly, we found TGF-β-repressors (*Smad7, Smurf2, Ski, Skil*) and genes involved in lymphocyte epithelial retention (*Itgae*) **(Fig. 7d, 7e, Supplementary information, Fig. 7c)**. Thus, SMAD4, directly by acting at the chromatin level could restrict TGF-β target gene expression. To identify putative partners of SMAD4 in WT and R2KO CD8 T cells, we conducted an enrichment motif analysis and uncovered similar motifs in the top 3 enriched motifs, notably ETS and RUNX family domains, indicating a potential interaction between SMAD4 and ETS or RUNX transcription factor families **(Fig. 7f, 7g)**. Thus, revealing that SMAD4 interacts likely with different partners to mediate its wide transcriptional impact in CD8 T cells.

Finally, to gain further insight into the epigenetic mechanisms by which SMAD4 mediates TGF-β target-gene repression before TGF-β-signal, we performed a ChIP of SMAD4 and a ChIP of a histone mark associated with gene expression, namely the acetylation of the 27^th^ lysine residue of the histone H3 protein (H3K27ac). We found that H3K27ac was enriched at the same SMAD4 binding regions of *Smad7, Skil* and *Itgae*, specifically in the absence of SMAD4 (SKO and R2SKO CD8 T cells). In contrast, we observed less enrichment in these SMAD4 binding regions in R2KO CD8 T cells. This indicates that SMAD4, in a TGF-β-independent manner and oppositely to TGF-β signal, promotes histone acetylation of TGF-β target gene promoters and enhancers mediating their repression **(Fig. 7h-i)**. Collectively, these findings highlight an upstream mechanism by which SMAD4, in CD8 T cells, mediates epigenetic control of a wide range of TGF-β target genes in the absence of TGF-β signaling, in anticipatory manner, and imposes a restriction of TGF-β signature, preventing IBDs.

## Discussion

Our study uncovers an uncharacterized critical feedforward regulation of the TGF-β effect governed by SMAD4 in CD8 T cells in a TGF-β independent manner, crucial to prevent chronic intestinal inflammation. Indeed, we reveal that the TGF-b-independent function of SMAD4 acts as a basal and active repressor of a myriad of TGF-β target genes, restraining the TGF-β signature in CD8 T cells in the absence of any TGF-β signaling. Therefore, ablation of SMAD4 impairs the effector predisposition of naïve CD8 T cells. However, SMAD4 ablation promotes CD8 T cell accumulation, and intestinal epithelial retention, in sharp contrast to total TGF-b signaling ablation. Besides, by inducing gene expression of the TGF-β negative feedback loop, SMAD4 ablation likely predisposes CD8 T cells to escape from the immune-regulatory effects of TGF-β and subsequently, combined with their incline for epithelial tropism, promotes their massive gut epithelial restricted chronic activation.

Although largely overlooked, emerging evidence over the last decade suggests that the presence and aberrant activation of CD8 T cells in the intestinal mucosa correlate with IBDs ^33,36,37^. Mechanisms leading to CD8 T cell activation in IBDs remain elusive. In a seminal work, Massague et al, demonstrated the importance of TGF-β in controlling T cell effector gene expression and suggested SMAD3 in the mechanism ^38^. However, the exact signaling branch of TGF-β, critical in establishing TGF-β-driven-immunosuppression, remains imprecise. We revealed that the absence of SMAD4 strongly impaired CD8 T cell response to TGF-β, thereby eliciting their activation that even a high dose of TGF-β or even a genetic activation of TGF-β remaining pathways cannot overcome. Our study demonstrated that amongst the different branches of TGF-β, SMAD4 appears to play a critical and non-redundant role in mediating immunosuppression induced by TGF-β in CD8 T cells.

Our current results reveal that TGF-β-independent SMAD4 function predisposes an effector differentiation program in naïve CD8 T cells. It has been described that SMAD4 contributes to T cell activation by inducing c-Myc during T cell activation ^22^. Here, we revealed that SMAD4-promoted T cell activation intervenes prior to any cognate antigen encounter. Our findings enforce studies questioning the previously presented naïve T cell dogma where naïve T cells are considered unpoised and homogeneous ^39^. Indeed, depending on the developmental origin, naïve CD8 T cells could be differentially “pre-programmed” thereby profoundly affecting their fate after peripheral cognate antigen encounter ^40^. However, the mechanism of this ‘pre-programing’ remains elusive. Our data suggest that, in contrast to TGF-β engagement, TGF-β-independent SMAD4 function promotes an effector commitment. Future investigations will be required to determine the molecular partners of SMAD4 that are important in mediating this effector engagement in naïve CD8 T cells.

SMAD4 deletion imposes a TGF-β-imprinting in CD8 T cells. TGF-β is critical for CD103 induction and IELs formation and retention in the gut ^34,41^. The fact that irradiated mice reconstituted with BM cells from TGF-βRII KO mice did not exhibit such a severe intestinal inflammation as TGF-βRII and SMAD4 double knockout mice is likely due to the lack of retention molecules such as CD103 in the former case. Indeed, ablation of SMAD4 in a setting of TGFβ-RII deficiency restores the epithelial retention capacity of CD8αβ T cells and may explain the more severe intestinal immunopathology observed in R2SKO compared to R2KO mice. Accordingly, CD103-blockade alleviates intestinal immunopathology in SMAD4-deficient mice. This indicates the crucial role exerted by SMAD4, in absence of TGF-β signal, in limiting epithelial retention of CD8 T cells.

Deletion of SMAD4, in some lymphocytes such as NK cells, enhances their response to TGF-β likely by curtailing the TGF-β-SMAD4-independent pathways (encompassing TRIM33 and the non-canonical pathways) ^42^. We have first envisaged a similar mechanism (of hyper-responsiveness to TGF-β) in our study. However, we unveiled that CD8 T cells lacking SMAD4 exhibit a strong defect to respond to TGF-β. Interestingly, this impressive defect was reminiscent of what is observed in patients suffering from IBDs and CRCs ^16^. Indeed, due to an elevated expression of the TGF-β repressor, SMAD7, T cells from those patients are not responsive to the immunoregulatory effect of TGF-β, highly enriched within the intestine ^3,17^. The impaired responsiveness to TGF-β explains why SMAD4 depletion could attenuate CD8 T cell effector differentiation on the one hand ^22^ but allows their chronic activation within the intestine, on the other hand, thereby reconciling this dichotomy. Indeed, insensitive to the TGF-β signal, SMAD4-deficient CD8 T cells could be chronically activated within the intestine, exhibit an effector program and contribute to IBDs.

SMAD4 deletion and total TGF-β signaling disruption have a striking opposite transcriptional and functional outcome. We show that SMAD4 binds to promoter regions of numerous TGF-β target genes and regulates inversely their expression in the absence of TGF-β signal by inducing epigenetic modifications such as chromatin acetylation. Indeed, before any TGF-β signal, SMAD4 restricts TGF-β signature, in an anticipatory mechanism, to potentiate and sensitize CD8 T cells to the effect of TGF-β once a TGF-β signal is received. Indeed this negative feedforward action limits the basal expression of TGF-β target genes and allow likely a better fine tune-regulation. In line with this concept of TGF-β -potentiation, by repressing a wide range of potent TGF-β repressors such as *Smad7, Ski, Skil* and *Smurf2*, SMAD4 facilitates TGF-β effect, in a TGF-β independent manner. This original negative feedforward regulation governed by SMAD4 explains the dual effect by which SMAD4 restrains TGF-β outcome, prior to TGF-β signal, but also potentiates it after TGF-βR engagement.

In summary, our study reveals that SMAD4 pre-conditions the fate of naïve CD8 T cells. We uncover a critical and non-redundant feedforward regulation governed by SMAD4 that finely preprograms naïve CD8 T cell homeostasis, with direct consequences on chronic intestinal inflammation.

## Supplementary figure legends

**Supplementary figure 1:**
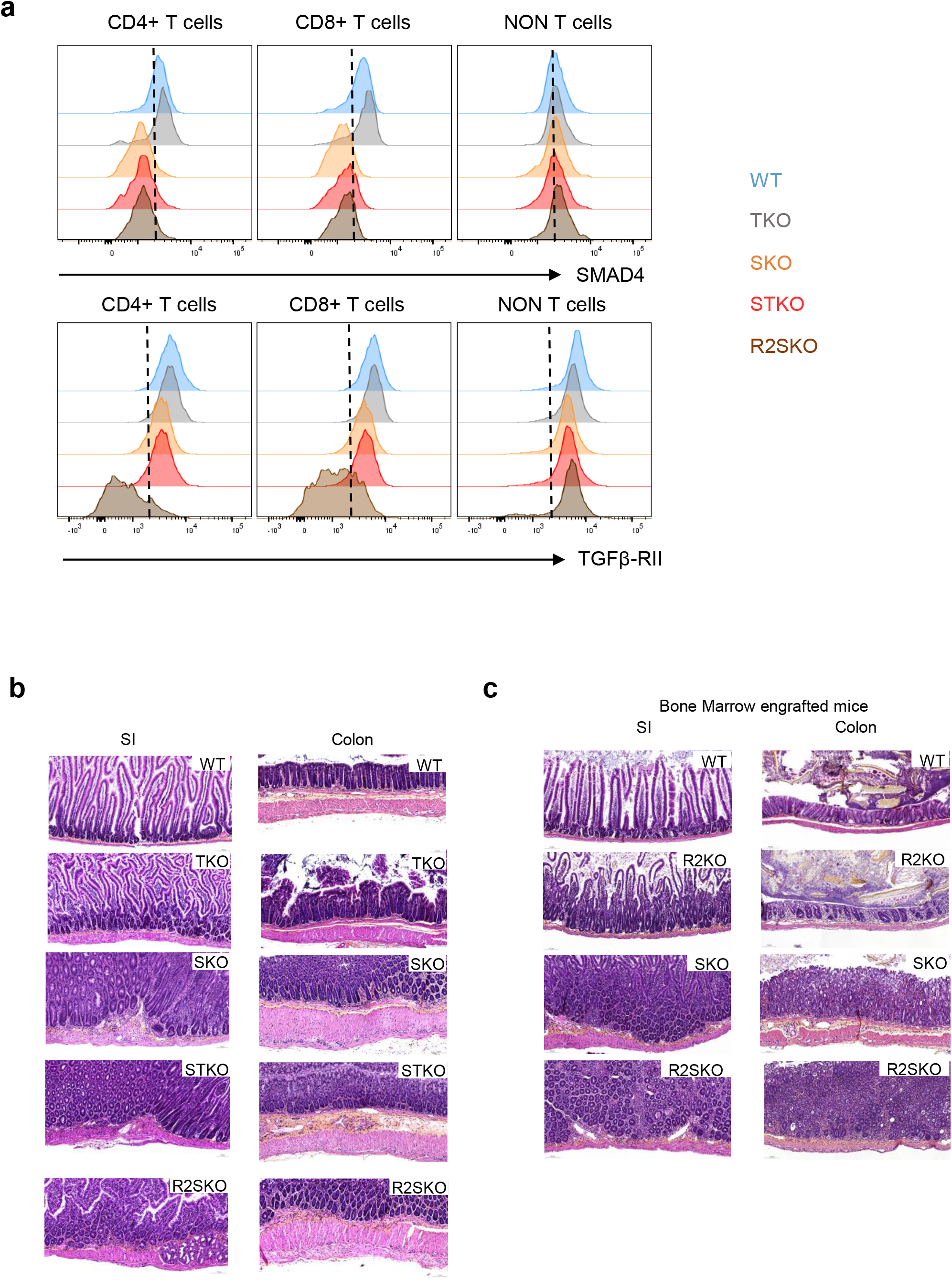
relative to figure 1. **(a):** Flow cytometry staining of SMAD4 and TGFβ-RII in T cells and non-T cells from WT, TKO, SKO, STKO and R2SKO mice**. (b):** Representative Hematoxylin & Eosin (H&E) staining of duodenum and colon sections from 7 months-aged WT, TKO, SKO, STKO and R2SKO mice. **(c):** Representative Hematoxylin & Eosin (H&E) staining of duodenum and colon sections of irradiated RAG2KO mice reconstituted with WT, R2KO, SKO, or R2SKO BM cells.

**Supplementary figure 2:**
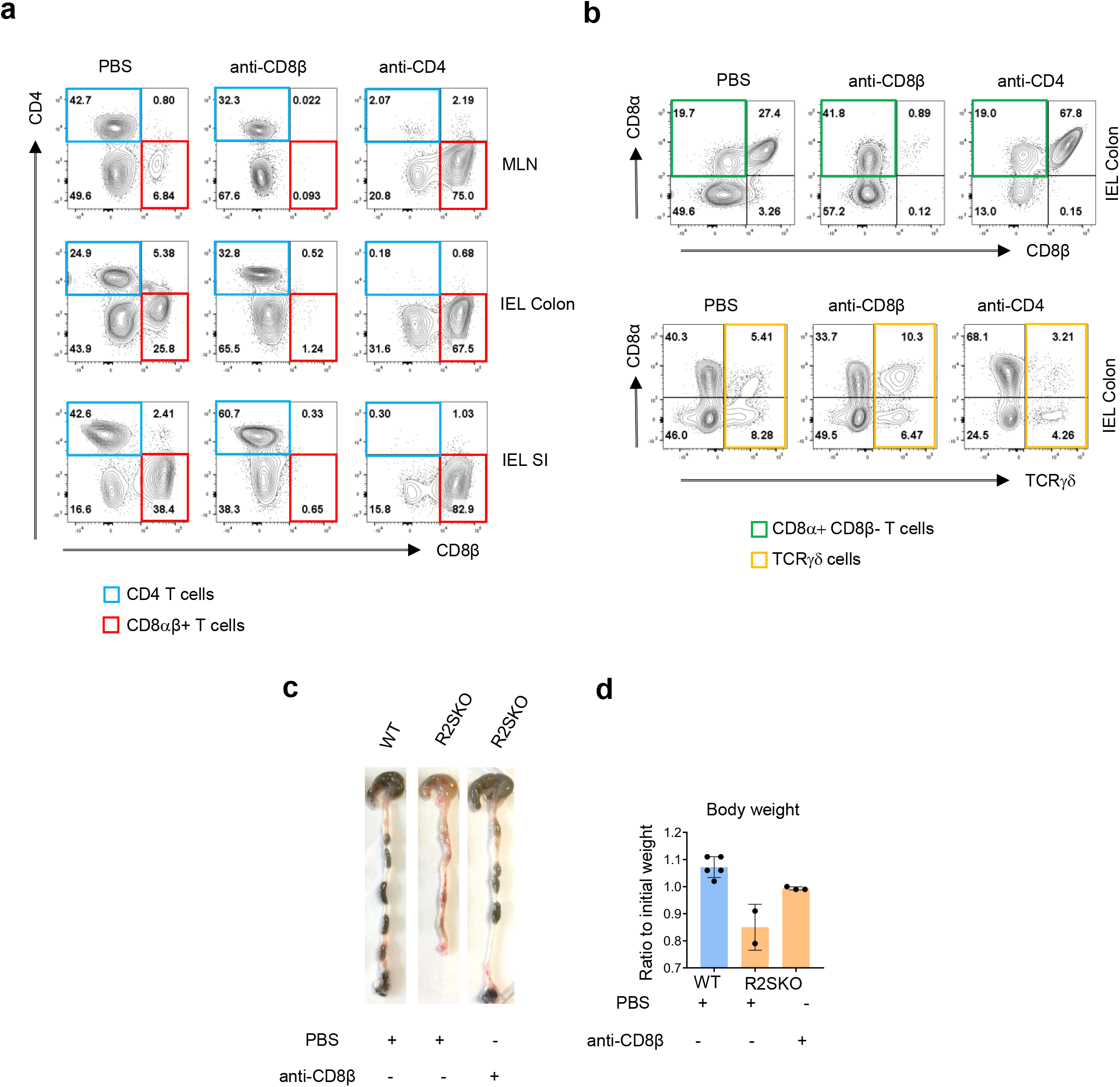
relative to figure 2. **(a-b):** Representative flow cytometry plots showing the frequency of CD4 and CD8αβ T cells **(a)**; TCRγδ and CD8αα **(b)** among CD45+ cells in the MLN and intra epithelial lymphocytes of irradiated RAG2KO mice reconstituted with WT BM cells and injected with PBS or anti-CD8β or anti-CD4 depleting antibody. **(c-d):** Representative pictures of colon **(c)** and body weight shown as relative to WT **(d)** of irradiated RAG2KO mice reconstituted with WT or R2SKO bone marrow cells and treated with anti-CD8β depleting antibody or PBS. **(e-g):** Experimental procedure for mixed bone marrow transplantation in irradiated RAG deficient mice **(e)**, colon length **(f)** and Body weight **(g)** of irradiated mice reconstituted with mixed WT and R2SKO BM cells. All data represent at least 3 independent experiments and presented as mean ± SD. Each symbol represents an individual mouse. Data were analyzed by unpaired Student t test. ns: nonsignificant; * p<0,05; **p < 0.01; ***p < 0.001; ****p < 0.0001.

**Supplementary figure 3:**
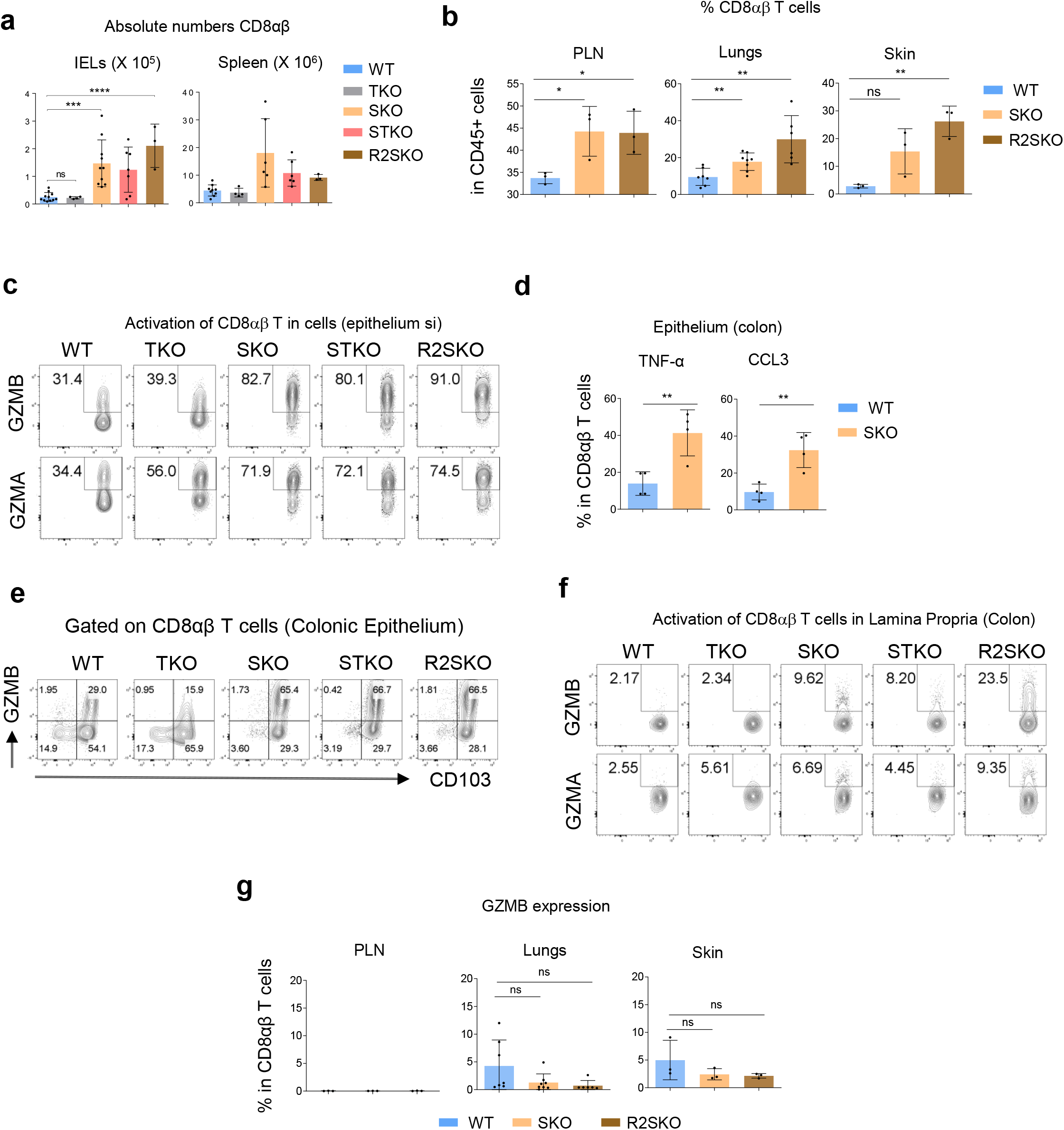
relative to figure 3. **(a):** Histograms showing the absolute numbers of CD8αβ T cells in the spleen and colonic intra epithelial lymphocytes of 7 months aged WT, TKO, SKO, STKO and R2SKO mice. **(b):** Histograms showing the frequency of CD8αβ T cells in the peripheral lymph nodes (PLN), lungs and the skin of 7 months-aged WT, SKO and R2SKO mice. **(c):** Flow cytometry staining of GZMA and GZMB in CD8αβ T cells of the epithelium of the small intestine of WT, TKO, SKO, STKO and R2SKO mice. **(d):** Histograms showing the frequency of TNF-α and CCL3 producing CD8αβ T cells in the colonic epithelium of 7 months-aged WT and SKO mice. **(e):** CD103 and GZMB staining showing that activated CD8αβ T cells are mainly CD103+ and present in the intestinal epithelium. **(f):** Flow cytometry staining of GZMA and GZMB in the colonic *lamina propria* CD8αβ T cells of WT, TKO, SKO, STKO and R2SKO mice. **(g):** The frequency of GZMB producing CD8αβ T cells in the PLN, lungs and the skin of 7 months-aged WT, SKO and R2SKO mice. All data represent at least 3 independent experiments and presented as mean ± SD. Each symbol represents an individual mouse. Data were analyzed by unpaired Student t test. ns: nonsignificant; * p<0,05; **p < 0.01; ***p < 0.001; ****p < 0.0001.

**Supplementary figure 4:**
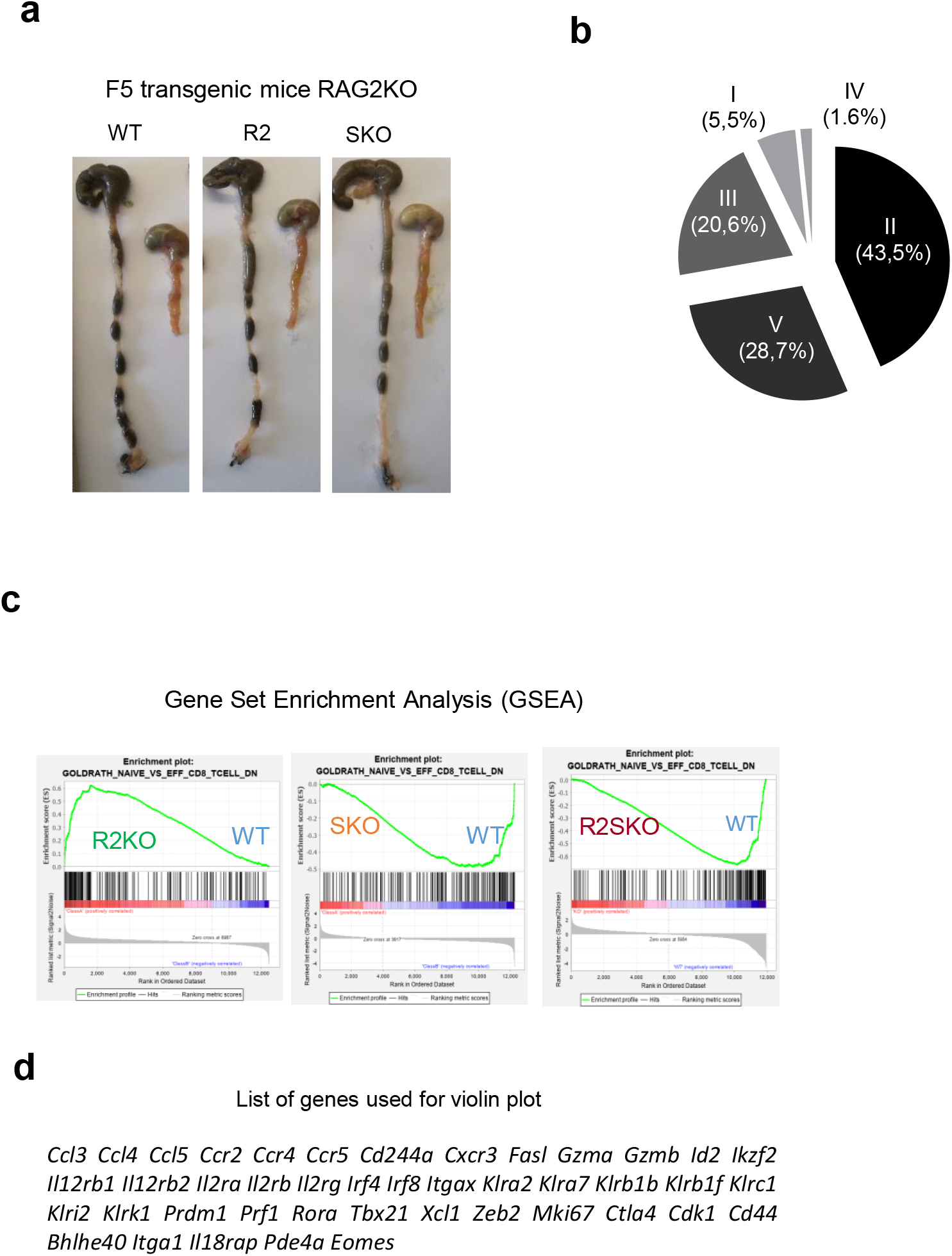
relative to figure 4. **(a):** Representative pictures of colon and duodenum from F5 TCR transgenic WT, R2KO and SKO 8 months-aged mice. **(b):** Pie chart showing the frequency of each cluster (related to Fig. 2b). **(c):** Gene Set Enrichment Analysis (GSEA) plot comparing gene expression arrays related to naïve or effector state of WT, R2KO, SKO, and R2SKO CD8αβ T cells. **(d):** List of the 43 selected genes related to the CD8 T cell effector state.

**Supplementary figure 5:**
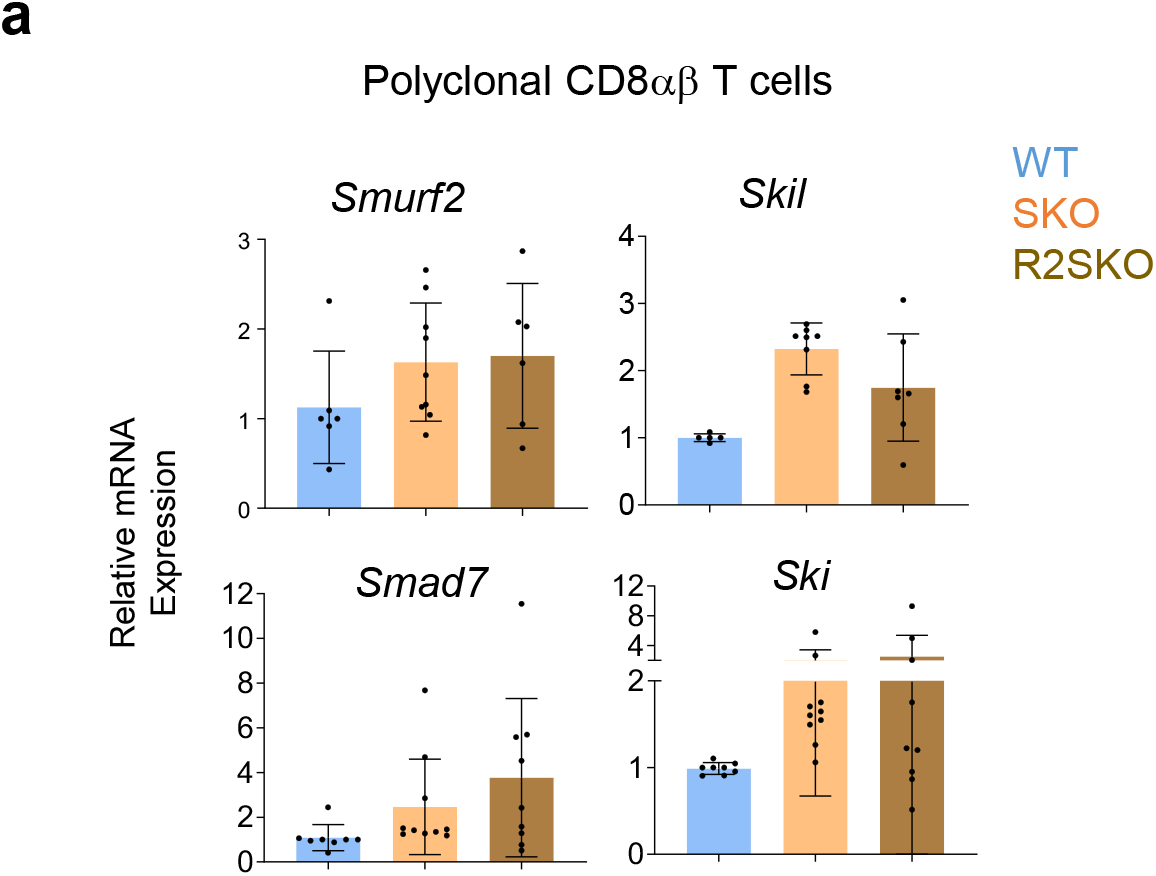
relative to figure 5. **(a):** Quantitative RT-PCR analysis of the expression of indicated TGF-β regulatory genes in polyclonal CD8αβ T cells from spleen of WT, SKO, and R2SKO mice. All data represent at least 3 independent experiments and presented as mean ± SD. Each symbol represents an individual mouse.

**Supplementary figure 6:**
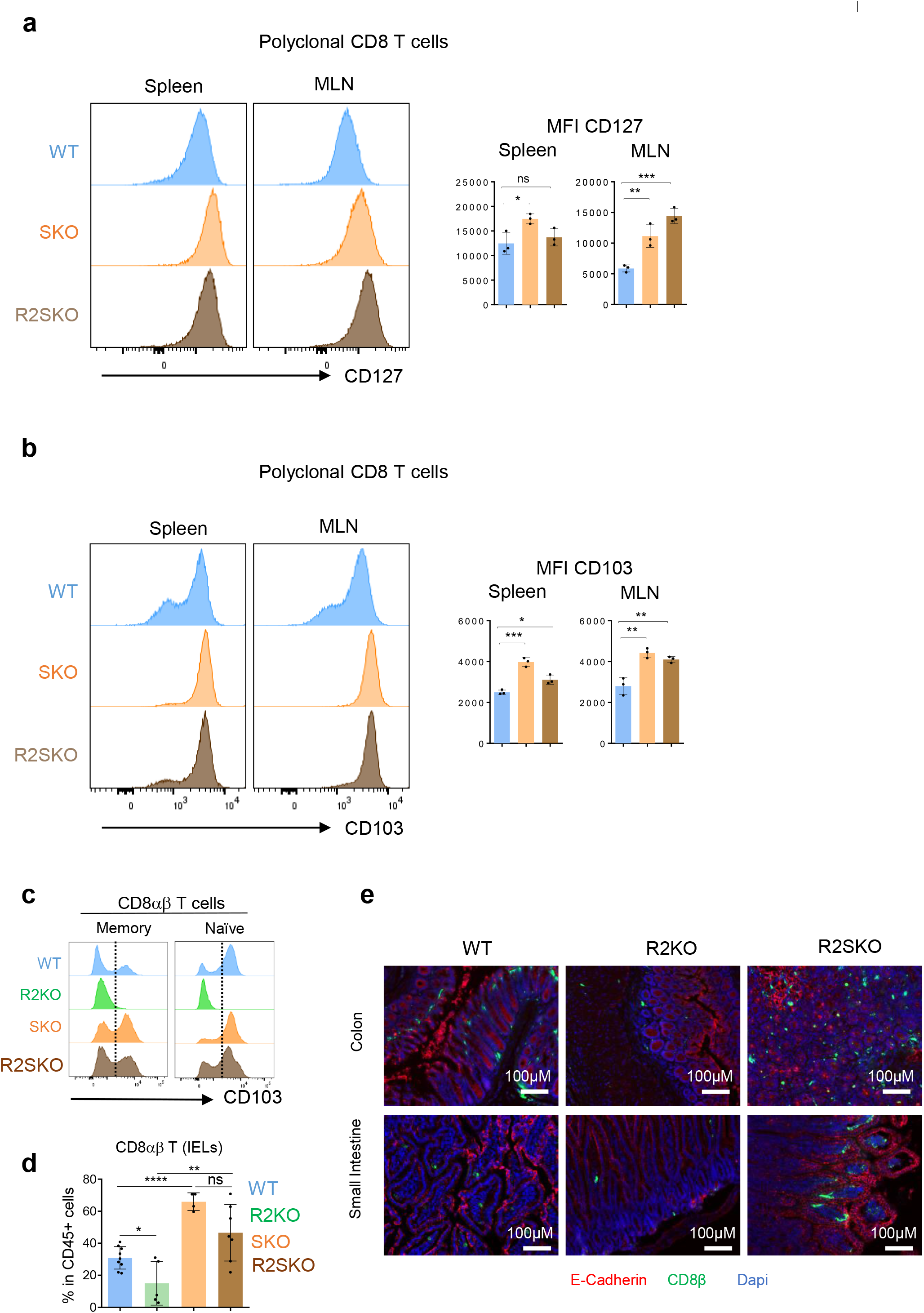
relative to figure 6. **(a-b):** Flow cytometry staining of CD127 **(a)** and CD103 **(b)** on polyclonal CD8αβ T cells in the spleen and MLN of WT, SKO and R2SKO mice. **(c):** Flow cytometry staining of CD103 on naïve (CD44 negative CD8 T cells) and memory (CD44 positive CD8 T cells) from irradiated and transplanted mice. **(d)** Histogram showing the frequency of CD8αβ T cells among CD45+ live cells in the colonic epithelium of RAG2KO irradiated mice and reconstituted with WT, R2KO, SKO or R2SKO BM cells **(e):** Representative pictures showing immune-fluorescence staining of CD8β (green), E-cadherin (red), DAPI (blue) in the small intestine and colon sections of RAG2KO irradiated mice reconstituted with WT, R2KO, or R2SKO BM cells. **(i):** All data represent at least 3 independent experiments and presented as mean ± SD. Each symbol represents an individual mouse. Data were analyzed by unpaired Student t test. ns: nonsignificant; * p<0,05; **p < 0.01; ***p < 0.001; ****p < 0.0001.

**Supplementary figure 7:**
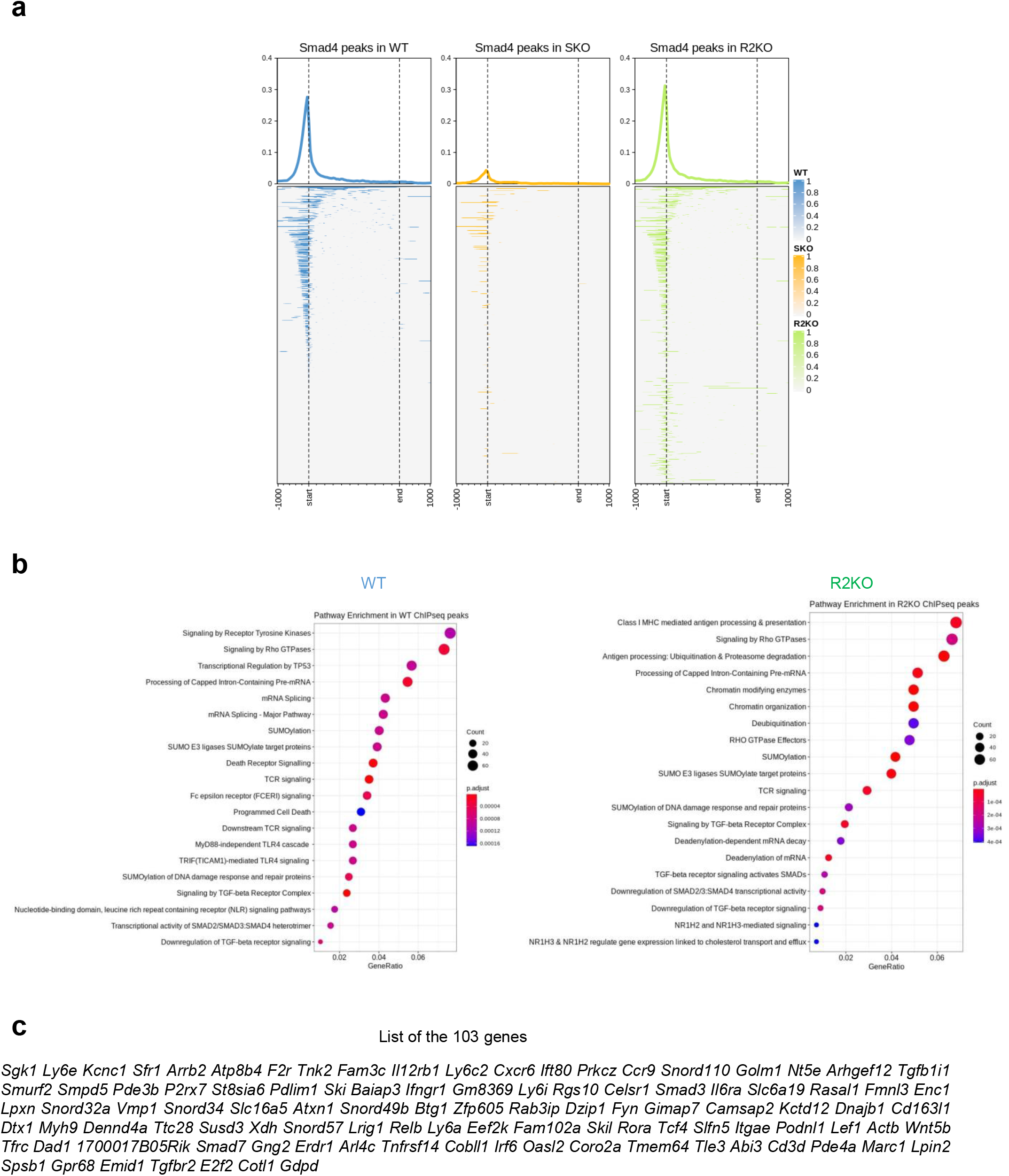
relative to figure 7. **(a)** Enriched heatmaps showing the SMAD4-occupancy signals in WT, SKO and R2KO (SKO was used as a negative control)**(a):** Biological pathway enrichment analysis in SMAD4 binding genes in WT and R2KO CD8 T cells **(b):** List of common genes that have a SMAD4 positive binding peak and are deferentially expressed between WT, SKO and R2KO naïve F5 CD8αβ T cells.

## Materiel and Method

### Mice

*CD4-Cre;Smad4^flox/flox^* (SKO), *CD4-Cre;Trim33^flox/flox^* (TKO)REF, *CD4-Cre;Smad^flox/flox^ Trim33^flox/flox^ (STKO), CD4-Cre;Smad^flox/flox^ Tgf-βRII^flox/flox^* (R2SKO), *CD4-Cre*; *Tgf-βRII^flox/flox^, CD4-Cre;Smad4^flox/flox^* stop^flox/flox^TGF-βRI^CA^ and RAG2KO mice were crossed and maintained in AniCan, a specific pathogen free animal facility of the Centre de Recherche en Cancérologie de Lyon (CRCL), Lyon, France. Unless mentioned otherwise, male and female mice were used. The experiments were performed in accordance with the animal care guidelines of the European Union and French laws and were validated by the local Animal Ethic Evaluation Committee (CECCAP).

### Antibiotic treatment

For antibiotics treatment, drinking water was supplemented with an antibiotic cocktail composed with Ampicillin (1g/L), Metronidazole (1g/L), Neomycin (1g/L), Vancomycin (0.5g/L) all purchased from Sigma-Aldrich. Antibiotic treatment was administrated just after weaning and until 5 months of age.

### Bone marrow transfer, CD8/CD4 depletion, and CD103 blockade

RAG2KO mice were irradiated (6 Gray) and reconstituted by intra-orbital injections with 10^6^ T cell-depleted bone marrow cells either from WT, R2KO, SKO or R2SKO mice. For the CD8/CD4 depletion, 20 days after BM reconstitution, mice received intraperitoneally 150μg of anti CD8β (clone. 53-5.8 Bioxcell) or anti CD4 (clone GK1.5, Bioxcell) once a week. We note that we used different antibody clones to verify the depletion by flow cytometry. For CD103 blockade, RAG2KO mice were irradiated (6 Gray) and reconstituted by intra-orbital injections with 10^6^ T cell-depleted bone marrow cells either from WT or SKO mice and 14 days after BM reconstitution, mice received intraperitoneally 100μg of anti CD103 blockade antibody (clone. M290 InVivoMab) or PBS 3 times per week.

### Histological Assessment of Inflammation

Colon and small intestine were fixed in 2% formaldehyde (VWR), embedded in paraffin and sectioned. Hematoxylin/eosin (Sigma Aldrich) staining was performed in embedded tissue. Intestinal inflammation was scored in a blinded fashion using a scoring system based on the following criteria: colon length, inflammatory cell infiltrate (severity and extent), crypt hyperplasia, presence of neutrophils within the crypts, presence of crypt abscesses, erosion, granulation tissues and villous blunting.

### Isolation of solenocytes, lymph nodes, lung, skin, intra epithelial and *lamina propria* cells

Spleens, peripheral (inguinal and axillary) or mesenteric lymph nodes were dissociated on nylon mesh and red blood cells were lysed with NH4Cl 9g/L (vol/vol). Lungs and ears were cut in small pieces and incubated in RPMI medium (Gibco) containing 20% Fetal Bovine Serum (Gibco), DNAse I Roche) at 100μg/ml and Collagenase from Clostridium Histolyticum (Sigma Aldrich) at 0.6mg/ml. For lungs, mice were perfused with PBS 1X, filtered and centrifuged on a percoll gradient 67%/44%. Small and large intestines were dissected after removing fat and payer patches. Intestines were longitudinally opened and washed in PBS 1X. Intestines were cut into small pieces and incubated with 5mM EDTA, 1mM DTT (Sigma Aldrich) at 37°C, under agitation. Epithelial cells and IELs were separated from tissue after 20 min. Tissues were then digested in RPMI medium (Gibco) containing 20% Fetal Bovine Serum (Gibco), DNAse I Roche) at 100μg/ml and Collagenase from Clostridium Histolyticum (Sigma Aldrich) at 0.6mg/ml. Intestinal LP was harvested from a 44% - 67% percoll gradient run for 20min at 1300 x g.

### Flow cytometry

Intra cellular and surface cell staining were performed using the following antibodies:

| Antibody | Fluorochrome | Clone | Company |
| --- | --- | --- | --- |
| CD8β | Alexa Fluor 488 | eBioH35-17.2 | Ebioscience |
| CD8β | Alexa Fluor 700 | YTS156.7.7 | Biolegend |
| CD8α | BV605 | 53-6.7 | Biolegend |
| CD45 | BV711 | 30-F11 | BD |
| CD3 | BV650 | 145-2C11/17A2 | BD/Biolegend |
| CD4 | V500 | RM4-5 | BD |
| Granzyme A | PerCP-eF710 | GzA 3G8.5 | eBiosciences |
| Granzyme B | APC | REA226 | Miltenyi |
| IFNγ | APC | XMG1.2 | eBiosciences |
| TNFα | Pe Cy7 | MP6-XT22 | Abcam |
| CCL3 | PE | REA355 | Miltenyi |
| CD103 | eF450/bv421 | 2E7/M290 | eBioscience/BD 678 |
| CD127 (IL-7Ra) | APC-eF780 | A7R34 | eBioscience 679 |
| CD44 | PerCP-Cy5.5 | IM7 | BD 680 |
| T-bet | PE-Cy7 | eBio4B10 | eBioscience 681 |
| Mouse anti mouse<br>E-cadherin | purified | 36/E-Cadherin | BD 682 |
| p-stat5 (p-Y694) | Alexa Fluor 647 | 47/Stat5 | BD 683 |
| Rabbit anti mouse<br>p-smad2<br>(Ser465/Ser467) | purified | E8F3R | Cell Signaling 684<br>685 |
| TCR$\gamma\delta$ | PE | GL3 | BD 686 |
| Donkey Anti mouse<br>IgG (H+L) | APC |  | Life Technologies 687<br>688 |
| Goat Anti rabbit<br>IgG (H+L) | A488 |  | Life Technologies 689<br>690 |

For IFNγ, TNFα, and CCL3 cytokine staining, cells were stimulated *ex vivo* for 4h with 1mg/ml PMA (Sigma Aldrich) and 1mg/ml Ionomycin (Sigma Aldrich) in presence of Brefeldin A (BD pharmingen) and Golgi Stop. After extracellular staining, cells were fixed and permeabilized using Cytofix/Cytoperm kit (BD) for IFNγ, TNFα, CCL3 staining and Fixation Permeabilization kit (invitrogen) for granzymes A/B and Tbet according to manufacturer’s protocol. For pSTAT5 and pSMAD2 intracellular staining, cells were fixed with 2% paraformaldehyde (EMS company) for 10 minutes at room temperature and then permeabilized with ice-cold Methanol for 30 minutes before intracellular staining. Flow cytometry data was acquired on BD LSR Fortessa using DIVA software and analysed by FlowJo software.

### Real time quantitative PCR

RNA was isolated with RNeasy mini kit (Qiagen) and reverse transcribed with iScript cDNA synthesis kit (Bio-rad). Real-time RT-PCR was performed using LightCycler 480 SYBR Green Master (Roche) and different set of primer (table) on LightCycler 480 Real-Time PCR System (Roche). Samples were normalized on GAPDH and analyzed according to the ΔΔCt method. There are the sequences of the primers used for qRT-PCR:

GAPDH: FW: 5’ CATGGCCTTCCGTGTTCCTA 3’ RV: 5’ TGTCATCATACTTGGCAGGT 3’
SMURF2: FW: 5’ AAACAGTTGCTTGGGAAGTCA3’ RV:5’ TGCTCAACACAGAAGGTATGGT3’
SKI: FW: 5’ TGACTCTGGACACAGCAGGA3’ RV:5’GAGAGGACAGCGAGGACAAG3’
SKIL: FW: 5’ AATAAAAAGCTGAACGGCATGGA3’ RV:5’GGGTTTTCCCATTGGCATGAAT3’
SMAD7: FW: 5’AAGTGTTCAGGTGGCCGGATCTCAG3’ RV: 5’ACAGCATCTGGACAGCCTGCAGTTG3’

### Bioinformatic Analyses

All genomic data was analysed with R/Bioconductor packages, R version 3.6.3 (2020-02-29) [https://cran.r-project.org/; http://www.bioconductor.org/].

### RNA-Seq

Illumina sequencing was performed on RNA extracted from triplicates of each condition. Standard Illumina bioinformatics analysis were used to generate fastq files, followed by quality assessment [MultiQC v1.7 https://multiqc.info/], trimming and demultiplexing. ‘Rsubread’ v1.34.6 was used for mapping to the hg38 genome and creating a matrix of RNA-Seq counts. Next, a DGElist object was created with the ‘edgeR’ package v3.26.7 [https://doi.org/10.1093/bioinformatics/btp616]. After normalization for composition bias, genewise exact tests were computed for differences in the means between groups, and differentially expressed genes (DEGs) were extracted based on an FDR-adjusted p value < 0.05 and a minimum absolute fold change of 2. DEG’ gene symbols were tested for the overlap with published signatures of interest using the ‘pathfindR’ package [https://doi.org/10.3389/fgene.2019.00858]. Hypergeometric Optimization of Motif EnRichment (HOMER v3.12) [https://doi.org/10.1016/j.molcel.2010.05.004] was used to calculate motif enrichment on the promoters of DEGs (up- and down-regulated genes separately), using default background settings.

### ChIP-Seq

ChIP libraries were prepared (Active Motif) and sequenced (Illumina NextSeq500) using a standard workflow. Resulting 75-nt single-end reads were mapped to the mm10 genome using the BWA algorithm with default settings [https://doi.org/10.1093/bioinformatics/btp324]. Only reads that passed Illumina’s purity filter, align with no more than 2 mismatches, and map uniquely to the genome were used in subsequent analyses. In addition, duplicate reads were removed. After normalization, the peak callers MACS/MACS2 [https://doi.org/10.1186/gb-2008-9-9-r137] were used to describe genomic regions with local enrichments in tag numbers relative to the Input data file (~ random background). Genomic ranges (‘GenomicRanges’ package) were used to perform genomic context annotations using the R packages ‘annotatr’ [DOI: 10.1093/bioinformatics/btx183], ‘ChIPSeeker’ [DOI: 10.1093/bioinformatics/btv145], and ‘ChipPeakAnno’ [DOI: 10.1186/1471-2105-11-237]. Enriched heatmaps (‘EnrichedHeatmap’) [https://doi.org/10.1186/s12864-018-4625-x] were used to visualize average ChIP peak signals. HOMER v3.12 was used to calculate motif enrichment in the vicinity of ChIP peaks.

All sequencing data has been uploaded into the GEO repository, with Accession number: XXXXXXXXXX

### qPCR-based ChIP

The PCR based ChIP was done using *ChIP It PBMC kit* from Active Motif catalog n° 53042 and ChIP It qPCR analysis kit from Active Motif catalog n° 53029. Cells were collected from freshly harvested spleen, MLN and peripheral lymph nodes and then sorted using CD8+ isolation kit from Miltenyi Biotec catalog n° 130-104-075. Chromatin preparation and immunoprecipitation (IP) were performed according to manufacturer’s protocol. IP were performed using anti-SMAD4 (EP618Y, Abcam); anti-H3K27ac (4729, Abcam); and rabbit IgG control (2729s, Cell Signaling). These are the sequences of the primers used por qPCR-based ChIP: SKIL promoter (SMAD4 binding site) FW: 5’ TATGACGGGCTAGCTTCACA 3’ RV: 5’ GAGACGGTAAGAGGTGGAGG 3’ CD103 (SMAD4 binding site) FW: 5’ggcagagcaaggatttgaac3’ RV: 5’CAGAGGCTcagagaaaatagcc3’ SMAD7 (SMAD4 binding site) FW: 5’ AAACCCGATCTGTTGTTTGC 3’ REV: 5’ GGCCGTCTAGACACCCTGT 3’

### In vitro survival assay and IL-7 response

CD8 T cells were obtained from freshly harvested mesenteric lymph nodes (MLN) from F5 transgenic WT, R2KO, SKO, and R2SKO mice. F5 naïve CD8 T cells were cultured in 96 well plate (10^5^ cells/well) in complete RPMI media with or without recombinant IL-7 at 10 ng/ml for different time points (0, 1, 3 and 5 days). For each time point, cells were washed and stained with LIVE/DEAD Fixable Dead Cell Stains kit (Life Technologies) according to manufacturer’s protocol, and fluorescent Abs against CD8 and CD45. The frequency of surviving CD8 T cells was determined by flow cytometry. For pSTAT5 staining, naïve CD8 T cells from the MLN of F5 transgenic WT, R2KO, SKO, and R2SKO mice were starved for 30 minutes at room temperature and then treated with mouse recombinant IL-7 at 10 ng/ml in RPMI 2% SVF for 30 minutes at 37°C. After 30 minutes of IL-7 stimulation, cells were immediately prepared for pSTAT5 staining (see Flow Cytometry section).

### *In vitro* CD8 T cells activation and differentiation

Briefly, splenic naïve CD8 T cells of WT, R2KO, and SKO mice were isolated using Mojosort negative selection kit from Biolegend and activated for 4 days via anti-CD3/anti-CD28 antibodies (10 μg/ml) plate bound (CD3, clone 145-2C11 catalog no 16-0031-86.; CD28, clone 37.51 catalog no. 16-0281-86). F5 naïve CD8 T cells were cultured in 96 well anti-CD3/anti-CD28 plate bound (10^5^ cells/well) in complete RPMI media with the presence of recombinant IL-7 at 10 ng/ml for all conditions. 4 days after, cells were washed and stained with LIVE/DEAD and Abs against CD45, CD8, Granzyme B and Tbet.

### *In vitro* TGF-β treatment and suppression assay

Splenic naïve CD8 T cells of WT, R2KO, and SKO mice were isolated using Miltenyi selection kit (Miltenyi Biotec) or Mojosort negative selection (Biolegend) and activated for 4 days via anti-CD3/anti-CD28 antibodies (as described above) in the presence or absence of human recombinant TGF-β1 (Miltenyi Biotec). The cells were cultured with TGF-β1 since the beginning. We note that we added IL-7 at 10 ng/ml for all our in vitro activation assays to maintain cells live. 4 days after, cells were washed and stained with LIVE/DEAD and Abs against CD45, CD8, Granzyme B and TBET. For p-SMAD2 staining, splenic naïve CD8 T cells from F5 transgenic WT, R2KO, SKO, and R2SKO mice were starved for 30 minutes at room temperature and then treated with human recombinant TGF-β1 at 10 ng/ml in RPMI 2% SVF for 20 minutes at 37°C. After 20 minutes of TGF-β1 stimulation, cells were immediately prepared for pSMAD2 staining (see Flow Cytometry section).

### Statistics

Unless mentioned otherwise two-tailed Student’s t test was used to calculate statistical significance. P values <0.05 were considered significant. ns: nonsignificant; *p < 0.05; **p < 0.01; p< 0.001; ****p < 0.0001. Statistics were performed using Prism Software.

## Author contributions

R. I and S.M.S planned, supervised and conducted the experiments. H.H, C.B, and A.D performed the bio-informatic analysis. R.I, H.H, S.M.S, N.B, D.B and J.C.M discussed the data and provided conceptual input. S.M.S and J.C.M provide financial resources. R.I and S.M.S wrote the manuscript.

## Acknowledgments

We thank the cytometry, histology, genomic and animal platforms of the CRCL and CLB for their help and assistance. In addition we thank Brigitte Manship, Anne-Marie Schmitt Verhulst and Gregoire Lauvau for discussing the paper.

